# Agentic-AI-ready genome-wide poxvirus–host interaction screen refined by a protein language model

**DOI:** 10.64898/2026.09.10.750412

**Authors:** Jacob Marcel Anter, Jason Mercer, Artur Yakimovich

## Abstract

Recent mpox outbreaks highlight the necessity to understand the interactions between poxviruses and the human host. These can be discovered systematically through screening for host genes involved in infection at the single-cell level using RNA interference. However, off-target effects and assay noise obscure true gene-phenotype relationships, hampering the discovery of therapeutically relevant targets. Here, we show that integrating protein-protein interaction information derived from a protein language model boosts the discovery of vaccinia virus-host interactions. We propose ICARus – a positive-unlabelled read-out refinement framework to achieve this. Our approach enhances the identification of human genes with potential antiviral function. We provide the raw and refined read-outs of a genome-wide screen for vaccinia virus host factors as an agentic-AI-enabled community resource. Our findings provide a generalisable strategy for robust hit prioritisation in functional screens, accelerating discovery across complex biological systems.

## Introduction

Poxviruses are large double-stranded DNA viruses that infect a broad range of human cells, establishing interactions with their host (1–4). Members of this family encode approximately 200 confirmed proteins (5) that modulate host signalling pathways, innate immune responses and cellular processes essential for viral replication and spread. Recent outbreaks of mpox have renewed attention to the public health relevance of poxviruses and underscore the need for a deeper mechanistic understanding of poxvirus-host interactions (6). Despite extensive study of vaccinia virus (VACV) as a prototypical poxvirus, many host factors that contribute to distinct stages of the viral life cycle remain incompletely characterised. This limited understanding of host dependencies constrains efforts to fully elucidate viral pathogenesis and to identify host-directed antiviral strategies.

RNA interference (RNAi) screening is an established experimental technique for the identification of host factors involved in viral infections. In this approach, genes are iteratively silenced, allowing for the interrogation of the role of individual genes in a biological process of interest (7, 8). However, RNAi is hampered by off-target effects (OTEs) involving binding of an siRNA molecule to an mRNA molecule other than its intended target (9, 10). Established avenues of addressing OTEs include, amongst others, chemical modification of RNAi reagents as well as in silico and statistical approaches (11–14). Even with advancements in RNAi technology, OTEs remain a persistent problem, obscuring true gene-phenotype relationships and thereby leading to false discoveries. Furthermore, the reproducibility of RNAi can be affected by experimental conditions, siRNA quality and cellular context. These factors collectively contribute to a partial or inconsistent mapping of host factors that modulate the life cycle of a virus of interest. Consequently, many host-pathogen interactions (HPIs) remain poorly characterised or lack functional validation.

Hit triage has benefited from the increasing availability of experimentally validated biological knowledge, with functional annotation data and protein-protein interaction (PPI) data as the prevailing data modalities (11–14). However, the coverage of public interaction databases remains inherently limited, motivating the development of artificial intelligence-based approaches able to generalise beyond experimentally validated PPI instances. Recent advances in PPI prediction could prove beneficial for this endeavour. Models such as SENSE-PPI (15), xCAPT5 (16) and MaTPIP (17) span diverse architectures while achieving strong predictive performance.

In this work, we performed an image-based, human genome-wide siRNA screen to identify host factors involved in VACV infection. To mitigate off-target effects and assay noise, we devised a previously unexplored PPI-guided hit triage strategy that improves VACV host factor prioritisation. This approach, named ICARus (Interactome-Corrected Analysis of RNAi screens), is a positive-unlabelled (PU) learning framework. It uses quantitative image features, as well as host and virus protein sequences, as input. ICARus computes PPI probabilities between human host proteins and VACV proteins, which serve as input for a PU learning model that enhances the recovery of established host factors while deprioritising biologically implausible hits. The PPI features, in turn, are derived from embeddings generated by a protein language model (pLM). Capturing sequence representations relevant to fundamental principles of protein biochemistry, the injection of this prior biological knowledge improves the prioritisation of VACV host factors compared to the uncorrected screen. We demonstrate that the injection of PPI-derived prior biological knowledge is a powerful new avenue for addressing screening assay noise and OTEs. Furthermore, we publish the raw and refined read-out obtained by our approach as a community resource termed ICARus-Pox (http://icaruspox.github.io). ICARusPox functions as an agentic harness, combining the screen dataset with domain-specific analytical functions and interfaces to large language models (LLMs). This enables AI-assisted, iterative interrogation of HPIs and supports the identification and prioritisation of candidate antiviral targets for further in vitro investigation.

## Results

### Genome-wide RNAi screen for human host genes involved in VACV infection

To systematically elucidate the yet unknown HPIs of VACV, we performed a genome-wide image-based RNAi screen (Fig. 1a). For this, we developed a VACV infection assay with a genetically engineered dual fluorescence reporter VACV expressing eGFP and mCherry upon infection from early and late VACV promoters, respectively. Assay readout was based on high-content fluorescence microscopy and a subsequent cell-segmentation-based image analysis pipeline yielding single-cell fluorescence readout upon siRNA gene knockdown (see Methods). Anti-eGFP and mock infection were used as positive and negative controls. However, upon inspection of the Z’-score (18) of the readouts, the best-performing positive and negative controls resulted in 0.43, suggesting the linear separability of the results was marginal. This likely stemmed from OTEs and assay noise obscuring true gene-phenotype relationships. Despite that, given the large size of the dataset, we hypothesised that the true gene-phenotype relationships may be obtained in a data-driven way. Specifically, we devised a strategy employing a pLM-based PPI prediction model to deconvolve the screen results through prioritising more likely HPIs and deprioritising less likely ones (Fig. 1b).

**Fig. 1.**
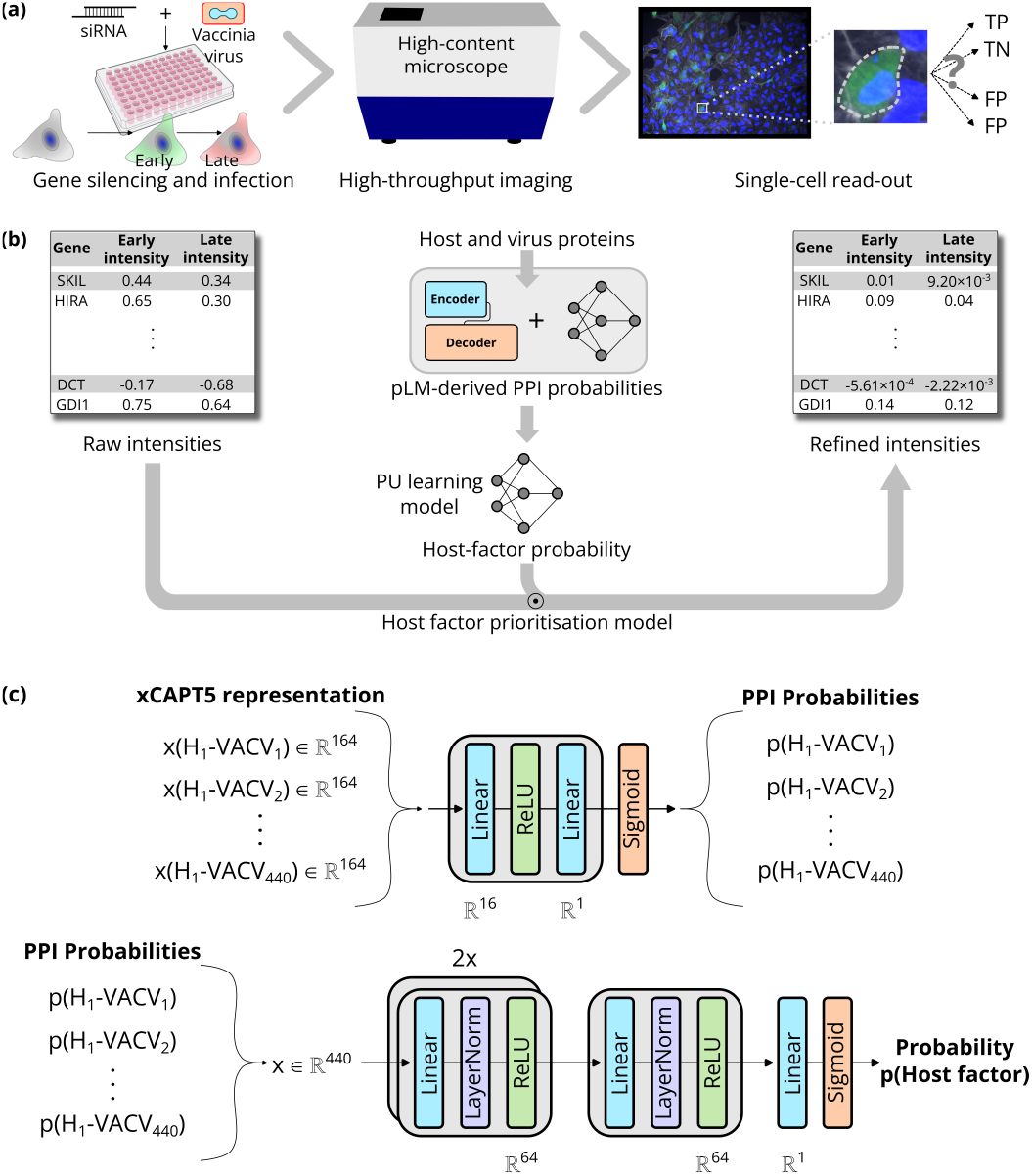
ICARus enables robust identification of VACV host factors: (a) Overview of the human genome-wide RNAi screen for identification of VACV host factors. (b) Overview of the read-out refinement pipeline, implementing a confidence-weighting (soft-masking) strategy. Prior biological knowledge is incorporated in two consecutive steps (PPI prediction followed by host-factor probability prediction) to compute a refinement factor for each gene. The screen read-out is multiplied by this refinement factor, weighting each gene according to its predicted likelihood of being a VACV host factor. (c) Detailed representation of the pipeline for predicting the probability that a gene is a VACV host factor. The pipeline comprises two sequential prediction stages. First, PPI probabilities are predicted between the human protein encoded by the gene of interest and all VACV proteins. These predicted PPI probabilities are then used as input to estimate the probability that the corresponding gene is a VACV host factor.

### Benchmarking protein-protein interaction models for VACV-human PPI prediction

To identify a suitable PPI prediction model for downstream modelling, we bench-marked three protein language model-based approaches for their ability to predict VACV-human interactions: xCAPT5 (16), SENSE-PPI (15) and MaTPIP (17). We first evaluated zero-shot performance on a curated VACV-specific PPI test set (see Methods). As shown in Table 1, xCAPT5 consistently outperformed the other models across multiple metrics. The best performance was achieved by xCAPT5 using the Pan checkpoint with an XGBoost classification head (AUROC = 0.81, PR AUC = 0.76, MCC = 0.54), whereas SENSE-PPI and MaTPIP showed lower overall predictive performance, particularly in terms of recall and MCC.

**Table 1.** Zero-shot performance of protein-protein interaction models on a VACV-human test set. MCC stands for Matthews Correlation Coefficient, AUROC stands for Area Under the Receiver Operating Characteristic Curve, and PR AUC stands for Area Under the Precision-Recall Curve. Best performance in bold.

| Model | Accuracy | Precision | Recall | F1-Score | Specificity | MCC | AUROC | PR AUC |
| --- | --- | --- | --- | --- | --- | --- | --- | --- |
| SENSE-PPI | 0.6143 | 0.7222 | 0.3714 | 0.4906 | 0.8571 | 0.2615 | 0.7527 | 0.7009 |
| xCAPt5 w/o XGBoost<br>Pan checkpoint | 0.7571 | 0.75 | **0.7714** | 0.7606 | 0.7429 | 0.5145 | 0.7857 | 0.6879 |
| xCAPt5 w/o XGBoost<br>Sled checkpoint | 0.5857 | 0.875 | 0.2 | 0.3256 | 0.9714 | 0.2694 | 0.6588 | 0.6448 |
| xCAPt5 w/ XGBoost<br>Pan checkpoint | **0.7714** | 0.7714 | **0.7714** | **0.7714** | 0.7714 | **0.5429** | **0.8090** | **0.7583** |
| xCAPt5 w/ XGBoost<br>Sled checkpoint | 0.5714 | 0.8571 | 0.1714 | 0.2857 | 0.9714 | 0.2381 | 0.6367 | 0.6590 |
| MaTPIP | 0.5571 | **1.0** | 0.1143 | 0.2051 | **1.0** | 0.2462 | 0.5404 | 0.6312 |

Based on this result, we selected xCAPT5 for further optimisation. To obtain more finely graded probability estimates suitable for downstream integration, we replaced the XGBoost classification head with a multilayer perceptron (MLP) (Fig. 1c) and fine-tuned the model on the VACV dataset. Fine-tuning was performed for both Sled and Pan checkpoints across five random initialisations each (Figs. S2-3). Performance metrics averaged across runs are summarised in Table 2.

**Table 2.** Performance of xCAPT5 after fine-tuning on VACV-human interaction data. Values represent mean ± s.d. across five random initialisations. Best performance in bold.

| Checkpoint | Accuracy | Precision | Recall | F1-Score | Specificity | MCC | AUROC | PR AUC |
| --- | --- | --- | --- | --- | --- | --- | --- | --- |
| Pan | 0.8057 | **0.8563** | 0.7371 | 0.7902 | **0.8743** | 0.6195 | 0.8635 | 0.8431 |
| | ± | ± | ± | ± | ± | ± | ± | ± |
|  | 0.0294 | **0.0358** | 0.0636 | 0.0367 | **0.0388** | 0.0568 | 0.0126 | 0.0081 |
| Sled | **0.8571** | 0.8211 | **0.9143** | **0.8648** | 0.8 | **0.72** | **0.9308** | **0.9364** |
| | ± | ± | ± | ± | ± | ± | ± | ± |
|  | **0.0256** | 0.0270 | **0.0361** | **0.0248** | 0.0361 | **0.0511** | **0.0117** | **0.0112** |

Despite its weaker zero-shot performance, the Sled check-point showed superior performance after fine-tuning, achieving higher AUROC (0.93 vs 0.86), PR AUC (0.94 vs 0.84) and MCC (0.72 vs 0.62) compared to the Pan checkpoint (Table 2). These results indicate improved generalisation of the Sled-based model in the supervised setting. Notably, the increased recall of the Sled-based model suggests enhanced sensitivity for capturing potential virus-host interactions, a key requirement for downstream host factor prioritisation. We therefore selected the Sled-based xCAPT5 model with an MLP classification head for integration into the downstream PU learning framework.

### A PU learning framework enables PPI-informed VACV host factor prioritization

To enable robust identification of VACV host factors, we developed a PU learning framework that leverages PPI-derived features (see Fig. 1b). The model was implemented as an MLP (Fig. 1c) trained with the non-negative PU (nnPU) loss (19) (see Methods). We also evaluated a multimodal variant combining phenotypic and PPI features, motivated by the hypothesis that both modalities provide complementary information. However, this approach did not outperform the PPI-only model (Fig. S7, Tables S3 and S5). This could suggest that the phenotypic features used here provide limited additional predictive information beyond the PPI-derived features, although more informative phenotypic representations may improve multimodal integration in future work. Additionally, this result may also be indicative of modality dominance or collapse. We therefore focused subsequent analyses on the PPI-only framework. We evaluated this framework under three training settings: uncorrupted PPI features, row-permuted PPI features, and column-permuted PPI features (see Methods, Figs. S6 and S8-9, Tables S4 and S6-7). As shown in Table 3, the model trained on uncorrupted PPI features achieved the best performance on the test set (AUROC = 0.77 ± 0.01, PR AUC = 0.45 ± 0.01).

**Table 3.**
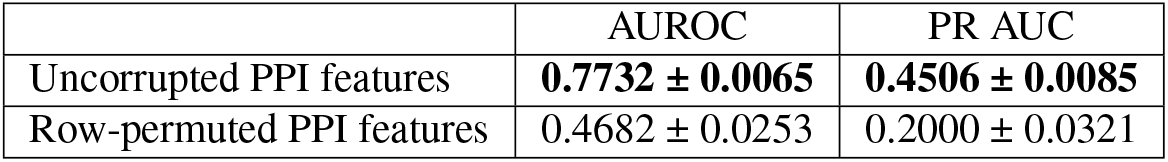
Performance of the PU learning framework under different training settings. Values represent mean ± s.d. across five random initialisations. Best performance in bold.

In contrast, the models trained on corrupted PPI features resulted in near-random performance (AUROC = 0.47 ± 0.03, PR AUC = 0.20 ± 0.03 for row permutation and AUROC = 0.50 ± 0.01, PR AUC = 0.23 ± 0.03 for column permutation), confirming that the PPI features contain meaningful predictive signal. Based on these results, the model using PPI features was used for downstream host factor prioritisation.

To validate model performance, we evaluated the model’s ability to discriminate known host factors from unlabelled genes using receiver operating characteristic (ROC) curves (Fig. 2a). Consistent with the metrics reported in Table 3, the model trained on uncorrupted PPI features (blue curve) achieves the highest AUC, whereas both models trained on corrupted PPI features follow the diagonal baseline and therefore perform close to random. Because functional screening data are highly imbalanced, ROC analysis alone does not fully capture the quality of the highest-ranked predictions. Precision-recall (PR) curves therefore complement the ROC analysis by quantifying the trade-off between precision and recall across different score thresholds (Fig. 2b). In line with the pattern exhibited by the ROC curves, the model trained on uncorrupted PPI features (PR AUC = 0.44) outperforms the models trained on corrupted PPI features (PR AUC = 0.22 for row-permuted PPI-features and PR AUC = 0.25 for column-permuted PPI features). The severe class imbalance notwithstanding, the model trained on uncorrupted PPI features maintains substantially higher precision across a broad range of recall values, indicating that highly ranked predictions are enriched for known host factors. Fig. 2c translates model performance into its practical impact on experimental validation by quantifying the enrichment of known host factors among the top-ranked predictions relative to random selection. The highest overall enrichment is achieved by the model trained on uncorrupted PPI features. Whereas the enrichment achieved by the corrupted models rapidly approaches the level expected from random selection, the enrichment of the uncorrupted model remains constantly elevated as the number of top-ranked genes varies widely. Collectively, these analyses demonstrate that our host factor prioritisation pipeline consistently assigns higher scores to known host factors than to unlabelled genes, which, in turn, translates into experimentally useful prioritisation. Because the evaluation treats all unlabelled genes as negatives, the reported performance metrics are conservative estimates. Some unlabelled genes are expected to represent undiscovered host factors and are therefore counted as false positives, implying that the true performance is likely to be underestimated.

**Fig. 2.**
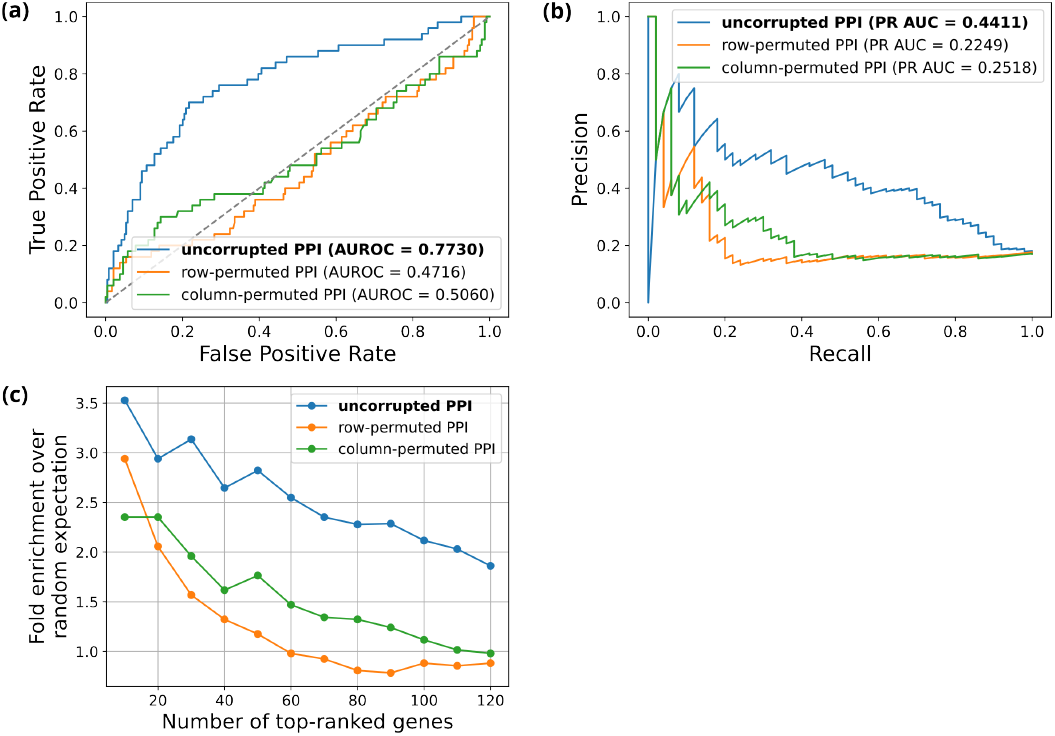
Visualisation of performance on the PU test set for representative runs. (a) ROC curves. The model trained on uncorrupted PPI features exhibits substantially improved discrimination between known host factors and unlabelled genes compared with models trained on corrupted features, which perform close to random. (b) Precision-recall (PR) curves. Despite the severe class imbalance, predictions made by the model trained on uncorrupted PPI features are substantially enriched for known host factors compared with models trained on corrupted features. (c) Fold enrichment of known host factors among the top-ranked predictions relative to random selection. The model trained on uncorrupted PPI features consistently achieves higher enrichment than models trained on corrupted features, indicating improved prioritisation for downstream experimental validation.

### PPI-informed confidence weighting broadens the functional landscape of top-ranked genes

The final step of our hit prioritisation pipeline involves soft-masking, i.e. multiplication of the raw intensities by the PU model probability (Fig. 1b). The soft-masking scales each intensity by the PU model-derived host-factor probability, thereby implementing a confidence-weighting strategy. Figs. 3a and 3b visualise the effect of this confidence weighting operation: In line with the PU learning prior of 0.05, the intensity of the vast majority of genes is pushed down towards zero. Genes assigned higher probabilities are consequently concentrated in the extreme tails.

**Fig. 3.**
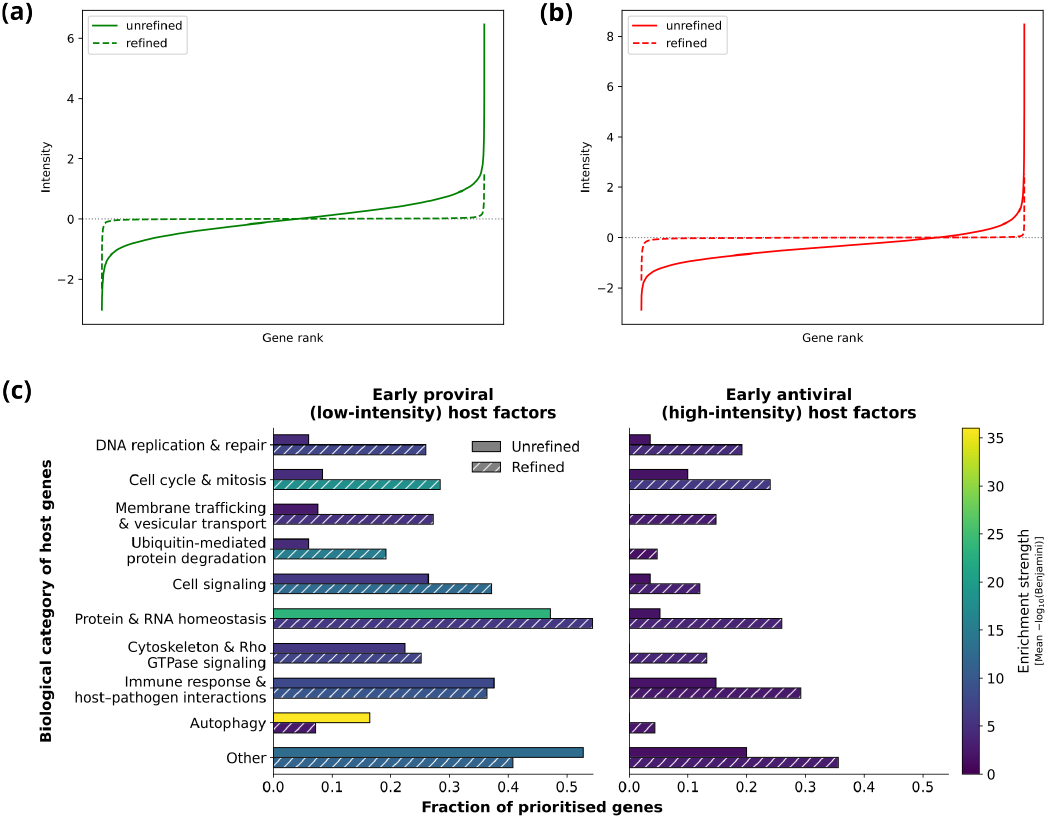
Effect of PPI-informed confidence weighting on host factor prioritisation. (a, b) Rank-intensity plots showing unrefined and refined early (a) and late (b) intensities. Refinement compresses the majority of intensities towards zero. (c) Effect of refinement on the functional landscape of top-ranked genes for early low (left) and early high (right) intensities. Horizontal bar length indicates the fraction of top-ranked genes assigned to each biological category, whereas bar colour indicates enrichment strength, quantified as the mean −log10(Benjamini) value of Reactome pathway terms assigned to the respective category. Refinement broadens the functional landscape of top-ranked genes and introduces functional structure into the high-intensity set.

To assess how this PPI-informed confidence weighting step alters host factor prioritisation, the top-ranked genes for the different intensity regimes were analysed via overrepresentation analysis (ORA) (see Methods, Table S10). Fig. 3c depicts the results for early low and high intensities; full results are depicted in Fig. S10. Fig. 3c reveals that PPI-informed confidence weighting broadens the functional landscape of the top-ranked genes, increasing the presence of previously weakly represented categories while retaining prominent functional categories. Noteworthy examples in the case of early low intensities include “Membrane trafficking & vesicular transport” and “Ubiquitin-mediated protein degradation”. For these categories, gene representation rose from 7.6% to 27.2% and 6.0% to 19.2%, respectively. For early high intensities, refinement expanded the gene representation of the “Immune response & host–pathogen interactions” category from 14.7% to 29.2%. Each of these categories encompasses processes with well-established roles in VACV infection. More general cellular processes also gain gene representation. Large shifts can be observed for “Cell signalling” (26.4% to 37.2%), “Cell cycle & mitosis” (8.4% to 28.4%) and “DNA replication & repair” (6.0% to 26.0%) in the early low intensities. The increased gene representation of these categories is consistent with established VACV biology: despite replicating entirely in the cytoplasm (20, 21), VACV interacts with host DNA damage response pathways and perturbs cell-cycle progression (22–24).

With regard to the early high intensities, it is striking that prior to refinement, the functional enrichment is remarkably sparse; several categories have no significant terms at all. However, refinement introduces additional functional structure into the high-intensity top-ranked set, with all categories gaining gene representation and categories already represented before refinement showing stronger enrichment. With regard to immune-related functions, the sparse enrichment may partly reflect the cellular context of the screen: In comparison to other cell types, HeLa cells mount a comparatively weak transcriptional response to VACV infection, with limited induction of interferon-stimulated genes (25). Because VACV actively antagonises interferon (IFN) production and signalling, depletion of host factors involved in these pathways may confer little additional advantage to the virus (26–28). Notably, VACV strain Western Reserve induces robust type I IFN signalling in bone marrow-derived macrophages, highlighting the cell-type dependence of this response (29). A second possibility is that antiviral host factors are often pleiotropic and also participate in essential housekeeping processes (30–33). Their depletion may therefore impair cellular fitness, indirectly reducing viral replication and masking an antiviral phenotype. Collectively, these results show that rather than merely reordering an unchanged functional landscape, PPI-informed confidence weighting alters the functional composition of top-ranked genes, broadening their representation across biologically coherent processes while retaining prominent functional categories.

### An interactive community resource for candidate antiviral target exploration

To facilitate access to the findings of this work for the virology community, we provide a publicly available online resource, ICARusPox (Fig. 4). Rather than merely serving as a static data repository or visualisation interface for the screen, ICARusPox provides an agentic harness in which the screen data can be interactively queried and analysed. Beyond browsing the screen results through the dashboard (Fig. 4a), the client-side JavaScript application allows users to execute scripts in a sandbox using screen-specific functions exposed through the BioJS namespace. Notably, users can connect to leading AI models via API (Fig. 4c), providing the AI model with the screen data and accompanying descriptions and instructions as context. This enables users to pose complex biological questions while leveraging the reasoning capabilities of LLMs. The context is further enriched through the inclusion of the latest Drug-Gene Interaction Database (DGIdb) (34), allowing users to explore putative antiviral drug candidates. Importantly, the agent can also generate and execute BioJS (35) code to interrogate the screen, enabling iterative, tool-assisted exploration of the data. By embedding an AI model within an interactive computational environment that provides both biological context and executable functionality, ICARusPox extends beyond a conventional question-answering system to provide an agentic framework for HPI discovery. ICARusPox can be accessed at icaruspox.github.io.

**Fig. 4.**
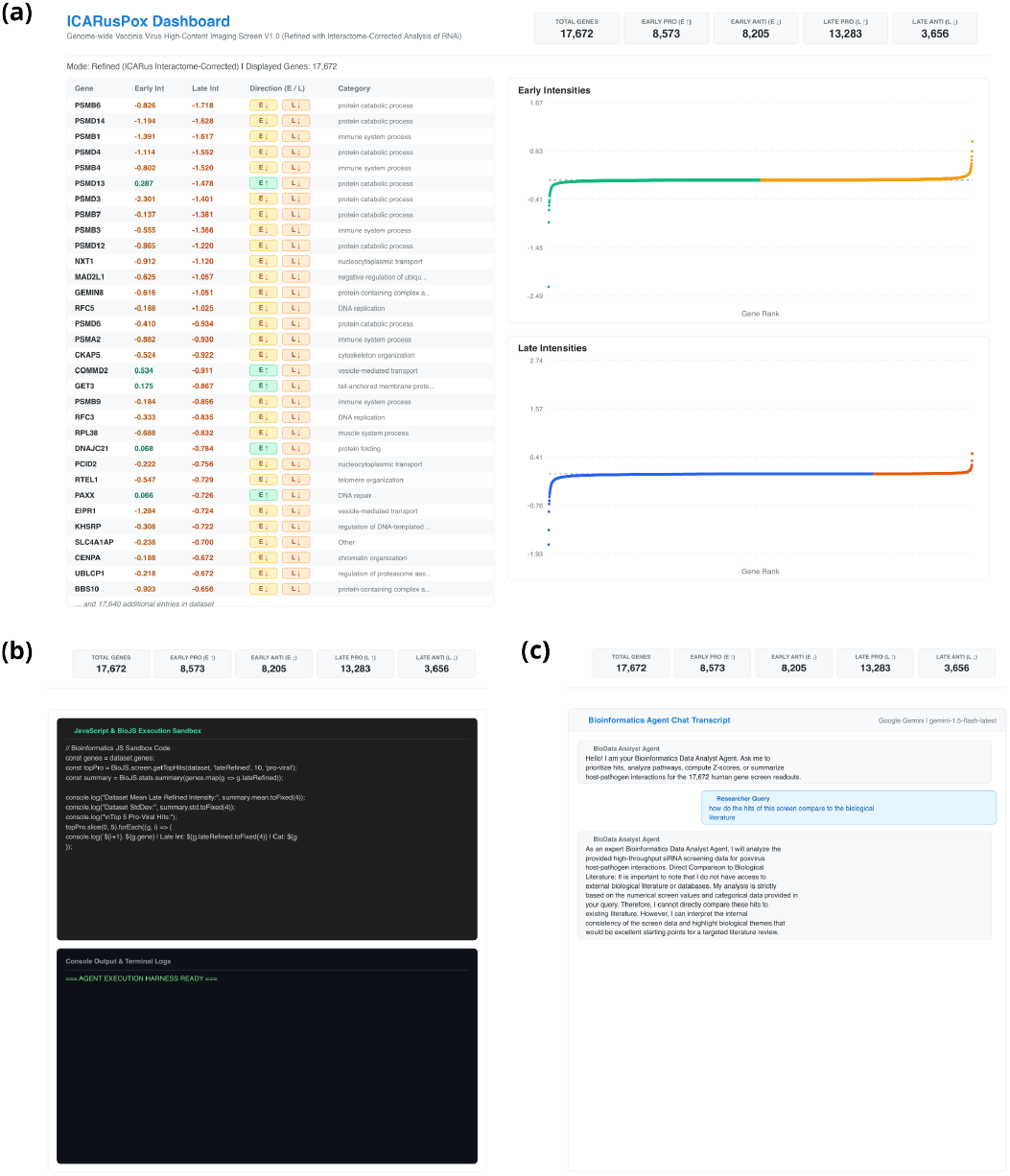
Interface of the ICARusPox online community resource. (a) The resource features a dashboard with the unrefined and refined intensities; the direction of the effect of the respective knockdown is also specified. Functional annotations provide further contextualisation. (b) A sandbox provides an isolated environment for executing BioJS(Gómez et al. 2013) code, allowing users to explore the screen without local installation. (c) Users can interact with the resource via an LLM of their choice, allowing in-depth analysis of the screen by e.g. targeted queries pertaining to a specific pathway, summarisation, etc.

## Discussion

In this work, we sought to make the analysis of an RNAi screen for host proteins interacting with VACV more robust to OTEs and assay noise. To this end, we developed ICARus, a positive-unlabelled learning framework capable of refining the read-out of a viral screen using PPI information and known host factors as prior biological knowledge. Applied to an image-based human genome-wide RNAi screen of VACV infection, ICARus enabled PPI-informed prioritisation of VACV host factors. We fine-tuned the xCAPT5 PPI prediction model on a VACV-specific PPI dataset and incorporated the resulting interaction probabilities into the PU learning framework. The resulting refinement altered the functional composition of the top-ranked genes, broadening their functional landscape in a fashion consistent with established aspects of VACV biology.

Finally, we make the resulting host factor prioritisation accessible through ICARusPox, providing a resource to support the selection of candidates for experimental validation. Beyond serving as a repository for the screen data, ICARusPox connects LLMs to the dataset and to domain-specific analytical functions, enabling AI-assisted exploration of HPI relationships. By coupling LLM-based reasoning to direct, structured access to the underlying data and executable analysis tools, the resource allows an LLM to iteratively formulate and test data-driven queries rather than merely generate responses based on a static representation of the screen. This agentic architecture therefore provides a means of translating open-ended biological questions into exploratory analyses, with the potential to accelerate hypothesis generation and the identification of candidate antiviral targets for experimental follow-up.

A central limitation of ICARus is its dependence on the quality and completeness of the biological prior used by the PU learning framework. Because the positive reference set comprises previously confirmed VACV host factors, the model may preferentially prioritise genes resembling known host factors while underprioritising host factors belonging to biological processes that are poorly represented in the reference set. The reduced representation of autophagy-related genes after refinement may illustrate this limitation. Evidence for the involvement of autophagy in VACV infection has accumulated in recent years (36–38), suggesting that the contribution of autophagy-associated host factors may be underestimated. The framework would therefore benefit from periodic curation and updating of the positive reference set as new experimental evidence becomes available.

A second limitation concerns the quality and biological context of the PPI data used for model fine-tuning. Experimentally demonstrated VACV-human PPIs may include interactions that are detectable under in vitro conditions but are unlikely to occur during infection, for instance because the interacting proteins are not co-localised in the relevant cellular compartments. Such interactions can introduce bias into the PPI-derived prior and consequently influence downstream hit prioritisation. Conversely, false-positive or low-confidence interactions arising from experimental errors may introduce noise into the prior. More extensive curation of the experimental confidence and cellular context of VACV-host interactions could therefore further improve the quality of the PPI-derived prior.

These limitations notwithstanding, ICARus represents, to our knowledge, the first application of PU learning to host factor prioritisation in viral RNAi screens. By treating genes not known to be host factors as unlabelled rather than negative instances, the framework explicitly accounts for the uncertainty inherent in biological discovery from such screens. Furthermore, using a PPI prediction model rather than a fixed interaction graph allows the biological prior to extend beyond experimentally established interactions. This may be particularly valuable for poorly characterised or newly emerging pathogens, for which experimentally mapped HPI networks are likely to be incomplete. The framework is also readily extensible to additional biological data modalities. Our attempt to integrate PPI-derived and phenotypic features showed, however, that multimodal integration does not inherently boost predictive performance, but that different modalities must provide complementary information. The aforementioned limitations, i.e. the dependence on the positive reference set as well as on the PPI data, are alleviated by the availability of both unrefined and refined intensities as the context of the agentic harness. This allows decisions to be made for each gene on a case-by-case basis, i.e. instead of dogmatically proposing genes associated with high refined scores as candidates, the agentic harness takes multiple aspects into account, thus permitting a more nuanced hit prioritisation.

ICARus and its future extensions may facilitate the study of poxvirus-host interactions and the identification of candidate antiviral targets. Although smallpox has been eradicated through globally coordinated efforts, the *Poxviridae* family continues to pose a public-health threat, as illustrated by the emergence of zoonotic mpox. The conservation of core biological processes and replication mechanisms among poxviruses suggests that VACV host factors identified by ICARus may provide a starting point for investigating host interactions in related poxviruses. Poxviruses are also valuable systems for dissecting HPIs and mechanisms of immune evasion, owing to their large genomes and extensive repertoire of host-modulating factors. In summary, ICARus provides a framework for prior-informed hit prioritisation in biological screens, with potential applications across pathogens and screening modalities.

## Code availability

Code developed in this study is available in the GitHub repository ICARusPox.

## Data availability

Data used in this study are available in the Zenodo repository at https://doi.org/10.5281/zenodo.22307677 (39).

## ACKNOWLEDGEMENTS

This work was partially funded by the Center for Advanced Systems Understanding (CASUS), which is financed by Germany’s Federal Ministry of Education and Research (BMBF) and by the Saxon Ministry for Science, Culture, and Tourism (SMWK) with tax funds on the basis of the budget approved by the Saxon State Parliament. A.Y. is supported by the Helmholtz Association Initiative and Networking Fund in the frame of Helmholtz AI as well as by the Helmholtz Foundation Model Initiative within the project “PROFOUND”. J.M.A. is supported by the CASUS Open Project.

## Methods

### VACV screening assay

The screening assay followed an identical protocol to (Rämö et al. 2014). HeLa CCL-2 cells obtained from ATCC were grown at 37°C under 5% CO2 in Dulbecco’s Modified Eagle Medium (DMEM, Invitrogen) containing 10% heat-inactivated fetal calf serum (Invitrogen). Automated liquid handling (BioTek EL406) was utilised for all stages of infection, fixation, and immunofluorescence. To initiate the assay, existing media in the RNAi-transfected plates was discarded and substituted with 40 µl of recombinant WR E EGFP/L mCherry VACV (Rämö et al. 2014) at a multiplicity of infection (MOI) of 0.125. Plates were maintained at 37°C for 1 hour to permit viral entry, after which the inoculum was swapped for 40 µl of DMEM supplemented with 10% FCS. At 8 hours post-infection (hpi), an additional 40 µl of DMEM/10% FCS containing 20 µM cytosine arabinoside (AraC) was introduced to inhibit secondary rounds of viral DNA replication. Cells were fixed at 24 hpi using 20 µl of 18% paraformaldehyde (PFA) for 30 minutes, followed by two 80 µl PBS washes. For EGFP detection, samples were incubated for 2 hours with a primary anti-GFP antibody (1:1000) diluted in 30 µl of a permeabilisation and blocking buffer (PBS containing 0.5% Triton X-100 and 0.5% BSA). Following two 80 µl PBS washes, cells were treated for 1 hour with 30 µl of secondary staining solution (PBS with 0.5% BSA) comprised of an Alexa Fluor 488-conjugated secondary antibody (1:1000), Hoechst dye (1:10000), and DY-647-phalloidin (1:1200; Dyomics). The procedure concluded with two final 80 µl PBS washes and the addition of 80 µl of H2O per well.

### Microscopy and image analysis

Automated image acquisition was carried out using Molecular Devices ImageXpress microscopes equipped with a 10X S Fluor objective (0.45 NA) and integrated with a Thermo Scientific robotic system for automated plate handling. Utilising the MetaXpress plate acquisition wizard, images were captured at a 12-bit dynamic range without gain across an equally spaced 3×3 grid (9 total sites per well) with overlap. The system employed laser-based focusing, with the site autofocus applied to all locations and the initial sample detection set to the first acquired well. While the Z-offset was determined manually, preliminary exposure times were established using the “AutoExpose” function and subsequently fine-tuned by hand to maximise the dynamic range while minimising overexposure. Fluorescence channels were selected based on the specific assay requirements. Following the acquisition, downstream image analysis and data normalisation were executed using adapted CellProfiler (Carpenter et al. 2006) pipelines identical to (Rämö et al. 2014) (Fig. S1, Table S1).

### Construction of the VACV-human PPI data set

The PPI dataset comprised experimentally validated positive interactions and computationally generated negative instances. Positive VACV-human PPIs were obtained from the Human-Virus Interaction Database (HVIDB) (40) and filtered for the VACV Western Reserve (WR) strain, resulting in 412 interactions after removal of deprecated entries.

Negative PPIs were constructed by pairing VACV proteins with human proteins localised to the nucleolus, leveraging the cytoplasmic replication of VACV, which makes stable interactions with nucleolar proteins unlikely (5, 20). Proteins known to interact with VACV were excluded from this set. To further reduce the likelihood of false negatives, candidate human proteins were required to share less than 50% sequence identity with any confirmed VACV interactor.

To minimise data leakage, train, validation and test splits were generated using a combination of sequence-based clustering and interaction network structure. Human nucleolar proteins were grouped by sequence similarity, whereas experimentally validated PPIs were represented as a bipartite graph and partitioned into connected components. Clusters and components were assigned exclusively to one of the three subsets.

Negative instances were generated separately for each subset by randomly pairing the corresponding human nucleolar and viral proteins, resulting in balanced datasets with a 1:1 ratio of positive to negative interactions. The final dataset comprised 686, 68 and 70 PPIs in the training, validation and test sets, respectively. Implementation details are provided in the Supplementary Information.

### Architecture and training of the MLP classification head for PPI prediction

The MLP classification head comprises two fully connected layers separated by a rectified linear unit (ReLU) activation. The first layer maps the xCAPT5-derived representations (ℝ^164^) to a 16-dimensional latent space, followed by a second layer projecting to a single logit (ℝ^1^). Logits are converted to probabilities using a sigmoid activation (Fig. 1c).

The model was implemented in PyTorch and trained for 100 epochs using the AdamW optimiser with a learning rate of 1 *×*10^*™*4^, weight decay of 1 *×* 10^*™*2^ and *ϵ* = 1*×*10^*™*8^. A dropout rate of 0.2 was applied after the ReLU activation. Training was performed with a batch size of 100 using binary cross-entropy loss.

### Construction of the PU learning data set

Starting from all genes interrogated in the RNAi screen, genes were filtered to retain only those associated with at least one encoded protein. Confirmed VACV host factors curated from published literature (Table S8) were used as positive instances, whereas all remaining genes were treated as unlabelled instances.

Phenotype feature vectors are three-dimensional vectors derived from the siRNA screen with the individual dimensions corresponding to standardised early viral gene expression, late viral gene expression and cell count (see Supplementary Information). These phenotype feature vectors were used to characterise gene-specific perturbation phenotypes. To preserve the distribution of phenotypic states across data splits, phenotype vectors were subjected to unsupervised clustering prior to splitting.

Because some genes encode multiple proteins and some proteins are shared across genes, splitting was performed at the level of connected components in a bipartite gene-protein graph. In this graph, genes were connected to all proteins encoded by the respective gene. Connected components were treated as indivisible units during splitting to prevent information leakage between training, validation and test sets arising from shared genes or proteins.

Connected components were assigned to training, validation and test sets using a balanced split strategy designed to approximately preserve predefined split proportions while distributing large connected components across splits. All splitting procedures were performed at the gene level to ensure consistency with the downstream prediction task.

To control class imbalance in the PU learning setting, a predefined positive-to-unlabelled (P:U) ratio was enforced independently within each split. Unlabelled genes were sampled in a phenotype-stratified manner to preserve the distribution of phenotype clusters across training, validation and test sets.

For each gene, protein-level VACV interaction probability vectors were aggregated into a single gene-level PPI feature vector by taking, for each dimension, the maximum value of the predicted interaction probability vectors across all proteins encoded by the respective gene. The resulting gene-level phenotype and PPI feature vectors were subsequently used as input to the PU learning architecture.

The final PU learning data set comprised 1.046, 104 and 294 genes in the training, validation and test sets, respectively. For the detailed composition of the PU learning data set, refer to Table S2. Implementation details are provided in the Supplementary Information.

### Architecture and training of the multimodal MLP for VACV host factor prediction

The PU learning framework was implemented as a multimodal multilayer perceptron (MLP) integrating phenotypic and PPI-derived features. The model comprises two parallel input branches for phenotypic and PPI features, respectively. Each branch consists of two processing blocks, each comprising a fully connected layer, layer normalisation, and a rectified linear unit (ReLU) activation (Fig. S4). The individual input branches were switchable, enabling training with both modalities or with either modality individually. The PPI-only architecture used for the final hit prioritisation is depicted in Fig. 1c.

Phenotypic input features were represented as three-dimensional vectors corresponding to standardised early viral gene expression, late viral gene expression and cell count. These features were projected to a 16-dimensional latent space.

PPI-derived features were constructed as 440-dimensional vectors, with each dimension corresponding to the predicted interaction probability between a human protein and one of the 440 VACV WR proteins. Interaction probabilities were obtained using xCAPT5 with the MLP classification head. For genes encoding multiple protein isoforms, feature vectors were aggregated by taking the element-wise maximum across all isoforms. The resulting feature vectors were projected to a 64-dimensional latent space.

For multimodal integration, sample-specific modality gating was employed (Fig. S5), whereby dedicated subnetworks learned the relative contribution of each modality for each individual sample rather than using global, fixed modality weights. The outputs of the subnetworks were fed into a softmax function to obtain modality weights, *α* for the phenotype modality and *β* for the PPI modality, satisfying *α* ∈ [0, 1], *β* ∈ [0, 1] and *α* + *β* = 1. The respective latent representations were multiplied by their corresponding modality weights and subsequently concatenated, resulting in an 80-dimensional joint representation. This representation was passed through a final processing block (fully connected layer, layer normalisation, and ReLU activation), followed by a final fully connected layer producing a single logit. Predicted probabilities were obtained by applying a sigmoid function to the logit. In the PPI-only setting, the 64-dimensional latent representation of the PPI branch was passed directly to the final processing block without modality gating.

The model was implemented in PyTorch and trained for 1,000 epochs using the Adam optimiser with a learning rate of 1 *×* 10^*™*3^, weight decay of 1 *×* 10^*™*4^ and *ϵ* = 1 *×* 10^*™*8^. Training was performed with a batch size of 32 using the non-negative PU (nnPU) loss (19) with a positive-class prior of *π*_*p*_ = 5 *×* 10^*™*2^.

### Functional landscape investigation of top-ranked genes via functional annotation clustering

To assess how the PPI-informed confidence weighting step depicted in Fig. 1b alters host factor prioritisation and whether this change is biologically reasonable, the top-ranked genes for the different intensity regimes were analysed. These different regimes are the following: early low, early high, late low and late high intensities for unrefined and refined intensities each. For each regime, the top 250 genes were selected, i.e. the 250 highest-ranked genes in the case of high intensities and the 250 lowest-ranked genes in the case of low intensities. These top 250 genes were subjected to an ORA using the functional annotation chart tool of the DAVID bioinformatics resources (41, 42) (Table S10). “REACTOME_PATHWAY” was used as the annotation category. As a background gene set, all genes interrogated in the screen have been taken. The EASE threshold has been set to 0.1 and the count threshold to 2. Significantly enriched Reactome pathway terms were selected using Benjamini < 0.05 as the selection criterion and assigned to ten broad biological categories. These are:

- Cell cycle & mitosis
- DNA replication & repair
- Cytoskeleton & Rho GTPase signalling
- Membrane trafficking & vesicular transport
- Immune response & host–pathogen interactions
- Protein & RNA homeostasis
- Ubiquitin-mediated protein degradation
- Autophagy
- Cell signaling
- Other

The precise mapping of Reactome pathway terms to these ten broad biological categories is listed in Table S9. For each combination of biological category and intensity regime, the average confidence (*™log*_10_(*Benjamini*) value) of the Reactome terms assigned to the respective biological category, as well as the fraction of the 250 top-ranked genes assigned to the respective biological category, were determined. These two quantities are visualised via a horizontal bar plot in Fig. 3c for early low and early high intensities. The results for all four regimes are displayed in Fig. S10. Note that one gene may participate in multiple biological categories, which is why the fractions do not necessarily sum up to 1.

### Evaluation metrics

The top k predictions refer to the k predictions with the highest probability. Recall@k is defined as the fraction of known hits recovered in the top k predictions (equation 1):

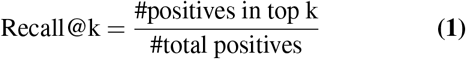

Precision@k is defined as the fraction of the top k predictions that are hits (equation 2):

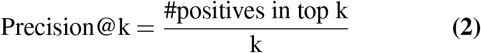

Enrichment@k is defined as the ratio between precision@k and the baseline positive rate (equation 3), where the baseline positive rate is defined as the fraction of total positives in the data set (equation 4).

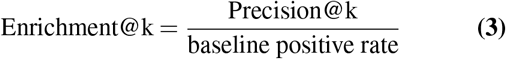

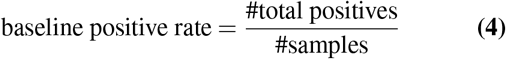

## Notes

https://icaruspox.github.io/

### Competing Interest Statement

The authors have declared no competing interest.

## References

1. Inger K Damon. Poxviruses. Manual of clinical microbiology, pages 1647–1658, 2011.

2. Sian Lant and Carlos Maluquer de Motes. Poxvirus interactions with the host ubiquitin system. Pathogens, 10(8):1034, 2021.

3. Bruno Hernaez and Antonio Alcamí. Poxvirus immune evasion. Annual Review of Immunology, 42(1):551–584, 2024.

4. Daniel Bourquain, Piotr Wojtek Dabrowski, and Andreas Nitsche. Comparison of host cell gene expression in cowpox, monkeypox or vaccinia virus-infected cells reveals virus-specific regulation of immune response genes. Virology journal, 10(1):61, 2013.

5. Stephen McCraith, Ted Holtzman, Bernard Moss, and Stanley Fields. Genome-wide analysis of vaccinia virus protein–protein interactions. Proceedings of the National Academy of Sciences, 97(9):4879–4884, 2000.

6. Rajesh Yadav, Anis Ahmad Chaudhary, Ujjwal Srivastava, Saurabh Gupta, Sarvesh Rustagi, Hassan Ahmed Rudayni, Vivek Kumar Kashyap, and Sanjay Kumar. Mpox 2022 to 2025 update: A comprehensive review on its complications, transmission, diagnosis, and treatment. Viruses, 17(6):753, 2025.

7. Christophe J Echeverri and Norbert Perrimon. High-throughput rnai screening in cultured cells: a user’s guide. Nature Reviews Genetics, 7(5):373–384, 2006.

8. Florian Heigwer, Fillip Port, and Michael Boutros. Rna interference (rnai) screening in drosophila. Genetics, 208(3):853–874, 2018.

9. Stephanie E Mohr, Jennifer A Smith, Caroline E Shamu, Ralph A Neumüller, and Norbert Perrimon. Rnai screening comes of age: improved techniques and complementary approaches. Nature reviews Molecular cell biology, 15(9):591–600, 2014.

10. Stephanie E Mohr and Norbert Perrimon. Rnai screening: new approaches, understandings, and organisms. Wiley Interdisciplinary Reviews: RNA, 3(2):145–158, 2012.

11. Jason Mercer, Berend Snijder, Raphael Sacher, Christine Burkard, Christopher Karl Ernst Bleck, Henning Stahlberg, Lucas Pelkmans, and Ari Helenius. Rnai screening reveals proteasome-and cullin3-dependent stages in vaccinia virus infection. Cell reports, 2(4): 1036–1047, 2012.

12. Li Wang, Zhidong Tu, and Fengzhu Sun. A network-based integrative approach to prioritize reliable hits from multiple genome-wide rnai screens in drosophila. BMC genomics, 10(1): 220, 2009.

13. Sandeep S Amberkar and Lars Kaderali. An integrative approach for a network based meta-analysis of viral rnai screens. Algorithms for Molecular Biology, 10(1):6, 2015.

14. Irene M Kaplow, Rohit Singh, Adam Friedman, Chris Bakal, Norbert Perrimon, and Bonnie Berger. Rnaicut: automated detection of significant genes from functional genomic screens. Nature methods, 6(7):476–477, 2009.

15. Konstantin Volzhenin, Lucie Bittner, and Alessandra Carbone. Sense-ppi reconstructs interactomes within, across, and between species at the genome scale. Iscience, 27(7), 2024.

16. Thanh Hai Dang and Tien Anh Vu. xcapt5: protein–protein interaction prediction using deep and wide multi-kernel pooling convolutional neural networks with protein language model. BMC bioinformatics, 25(1):106, 2024.

17. Shubhrangshu Ghosh and Pralay Mitra. Matpip: A deep-learning architecture with explainable ai for sequence-driven, feature mixed protein-protein interaction prediction. Computer Methods and Programs in Biomedicine, 244:107955, 2024.

18. Ji-Hu Zhang, Thomas DY Chung, and Kevin R Oldenburg. A simple statistical parameter for use in evaluation and validation of high throughput screening assays. Journal of biomolecular screening, 4(2):67–73, 1999.

19. Ryuichi Kiryo, Gang Niu, Marthinus C Du Plessis, and Masashi Sugiyama. Positive-unlabeled learning with non-negative risk estimator. Advances in neural information processing systems, 30, 2017.

20. Frank S De Silva and Bernard Moss. Origin-independent plasmid replication occurs in vaccinia virus cytoplasmic factories and requires all five known poxvirus replication factors. Virology journal, 2(1):23, 2005.

21. Utz Fischer, Julia Bartuli, and Clemens Grimm. Structure and function of the poxvirus transcription machinery. In The enzymes, volume 50, pages 1–20. Elsevier, 2021.

22. Caroline K Martin, Jerzy Samolej, Annabel T Olson, Cosetta Bertoli, Matthew S Wiebe, Robertus AM De Bruin, and Jason Mercer. Vaccinia virus arrests and shifts the cell cycle. Viruses, 14(2):431, 2022.

23. Matthew D Weitzman and Amélie Fradet-Turcotte. Virus dna replication and the host dna damage response. Annual review of virology, 5(1):141–164, 2018.

24. Antonio Postigo, Amy E Ramsden, Michael Howell, and Michael Way. Cytoplasmic atr activation promotes vaccinia virus genome replication. Cell reports, 19(5):1022–1032, 2017.

25. Kathleen H Rubins, Lisa E Hensley, David A Relman, and Patrick O Brown. Stunned silence: gene expression programs in human cells infected with monkeypox or vaccinia virus. PloS one, 6(1):e15615, 2011.

26. Geoffrey L Smith, Camilla TO Benfield, Carlos Maluquer de Motes, Michela Mazzon, Stuart WJ Ember, Brian J Ferguson, and Rebecca P Sumner. Vaccinia virus immune evasion: mechanisms, virulence and immunogenicity. Journal of General Virology, 94(11):2367–2392, 2013.

27. Beatriz Perdiguero and Mariano Esteban. The interferon system and vaccinia virus evasion mechanisms. Journal of Interferon & Cytokine Research, 29(9):581–598, 2009.

28. Jennifer H Stuart, Rebecca P Sumner, Yongxu Lu, Joseph S Snowden, and Geoffrey L Smith. Vaccinia virus protein c6 inhibits type i ifn signalling in the nucleus and binds to the transactivation domain of stat2. PLoS pathogens, 12(12):e1005955, 2016.

29. Siti Khadijah Kasani, Huei-Yin Cheng, Kun-Hai Yeh, Shu-Jung Chang, Paul Wei-Che Hsu, Shu-Yun Tung, Chung-Tiang Liang, and Wen Chang. Differential innate immune signaling in macrophages by wild-type vaccinia mature virus and a mutant virus with a deletion of the a26 protein. Journal of virology, 91(18):10–1128, 2017.

30. Meilin Li, Dingkun Peng, Hongwei Cao, Xiaoke Yang, Su Li, Hua-Ji Qiu, and Lian-Feng Li. The host cytoskeleton functions as a pleiotropic scaffold: orchestrating regulation of the viral life cycle and mediating host antiviral innate immune responses. Viruses, 15(6):1354, 2023.

31. Junru Yang, Ying Qu, Zhixiang Yuan, Yufei Lun, Jingyu Kuang, Tong Shao, Yanhua Qi, Yingying Li, and Lvyun Zhu. Targeting host dependency factors: A paradigm shift in antiviral strategy against rna viruses. International journal of molecular sciences, 27(1):147, 2025.

32. Masmudur M Rahman, Ami D Gutierrez-Jensen, Honor L Glenn, Mario Abrantes, Nissin Moussatche, and Grant McFadden. Rna helicase a/dhx9 forms unique cytoplasmic antiviral granules that restrict oncolytic myxoma virus replication in human cancer cells. Journal of virology, 95(14):10–1128, 2021.

33. Paul A Rowley, Brandon Ho, Sarah Bushong, Arlen Johnson, and Sara L Sawyer. Xrn1 is a species-specific virus restriction factor in yeasts. PLoS Pathogens, 12(10):e1005890, 2016.

34. Matthew Cannon, James Stevenson, Kathryn Stahl, Rohit Basu, Adam Coffman, Susanna Kiwala, Joshua F McMichael, Kori Kuzma, Dorian Morrissey, Kelsy Cotto, et al. Dgidb 5.0: rebuilding the drug–gene interaction database for precision medicine and drug discovery platforms. Nucleic acids research, 52(D1):D1227–D1235, 2024.

35. John Gómez, Leyla J García, Gustavo A Salazar, Jose Villaveces, Swanand Gore, Alexander García, Maria J Martín, Guillaume Launay, Rafael Alcántara, Noemi Del-Toro, et al. Biojs: an open source javascript framework for biological data visualization. Bioinformatics, 29(8):1103–1104, 2013.

36. Melanie Krause, Jerzy Samolej, Artur Yakimovich, Janos Kriston-Vizi, Moona Huttunen, Samuel Lara-Reyna, Eva-Maria Frickel, and Jason Mercer. Vaccinia virus subverts xenophagy through phosphorylation and nuclear targeting of p62. Journal of Cell Biology, 223(6):e202104129, 2024.

37. Yongge Li, Xu Miao, Rui Jia, and Ruikang Liu. Emerging interplays between poxviruses and autophagy. Frontiers in Cellular and Infection Microbiology, 15:1662511, 2025.

38. Melanie Krause, Artur Yakimovich, Noemi Vágó, Ingo Drexler, and Jason Mercer. Granularity screening identifies candidate genes involved in vaccinia virus induced lc3 lipidation. bioRxiv, pages 2026–03, 2026.

39. PhD Artur Yakimovich. Icaruspox/icaruspox.github.io: 0.0.1, September 2026.

40. Xiaodi Yang, Xianyi Lian, Chen Fu, Stefan Wuchty, Shiping Yang, and Ziding Zhang. Hvidb: a comprehensive database for human–virus protein–protein interactions. Briefings in bioinformatics, 22(2):832–844, 2021.

41. Da Wei Huang, Brad T Sherman, and Richard A Lempicki. Bioinformatics enrichment tools: paths toward the comprehensive functional analysis of large gene lists. Nucleic acids research, 37(1):1–13, 2009.

42. Da Wei Huang, Brad T Sherman, and Richard A Lempicki. Systematic and integrative analysis of large gene lists using david bioinformatics resources. Nature protocols, 4(1): 44–57, 2009.

