## Supplemental Information for "Agentic-AI-ready genome-wide poxvirus–host interaction screen refined by a protein language model"

### Supplementary information to the manuscript: Agentic-AI-ready genome-wide poxvirus–host interaction screen refined by a protein language model

**Authors**: Jacob Marcel Anter, Jason Mercer, Artur Yakimovich

#### Alleviating plate-to-plate variation via control-based Z-scoring

To mitigate plate-to-plate variation, the phenotypic features measured in the screen - early viral gene expression (eGFP fluorescence intensity), late viral gene expression (mCherry fluorescence intensity) and cell count - were standardised using control-based Z-scoring on a per-plate basis according to equation 1:

$$\mathrm{z} = \frac{\mathrm{x} - \mu}{\sigma} \tag{1}$$

where x denotes the raw measurement, $\mu$ the mean, $\sigma$ the standard deviation and z the resulting z-score. For each plate, $\mu$ and $\sigma$ were calculated exclusively from the corresponding control wells.

Control-based rather than conventional Z-scoring was employed because, although all plates contained identical controls, the remaining plate content differed between plates. Such differences in plate composition can introduce systematic biases and cause genes with similar phenotypic effects to yield substantially different Z-scores across plates. By computing $\mu$ and $\sigma$ solely from control wells, the composition of which was identical across all plates, plate-specific composition biases were minimised, and all plates were anchored to a common baseline, thereby enabling direct cross-plate comparison of Z-scores.

For both early and late viral gene expression, fluorescence intensities were quantified for four segmentation-defined compartments: whole-cell, nuclear, perinuclear and Voronoi-defined cellular regions. Together with cell count, this resulted in a total of nine readouts (four readouts for early viral gene expression, four readouts for late viral gene expression and cell count).

Control-based Z-scoring was performed independently for each of the nine readouts. Specifically, for each readout, plate-specific $\mu$ and $\sigma$ values were determined from the corresponding control wells and all raw measurements were standardised according to equation 1.

The effectiveness of control-based Z-scoring in alleviating plate-to-plate variation was assessed via UMAP dimensionality reduction followed by visualisation of individual measurements, where each measurement consisted of all nine readouts. Pronounced plate-specific clustering indicates the presence of batch effects, whereas a more homogeneous distribution indicates successful mitigation thereof. UMAP scatter plots for selected controls before and after control-based Z-scoring are shown in Fig. S1.

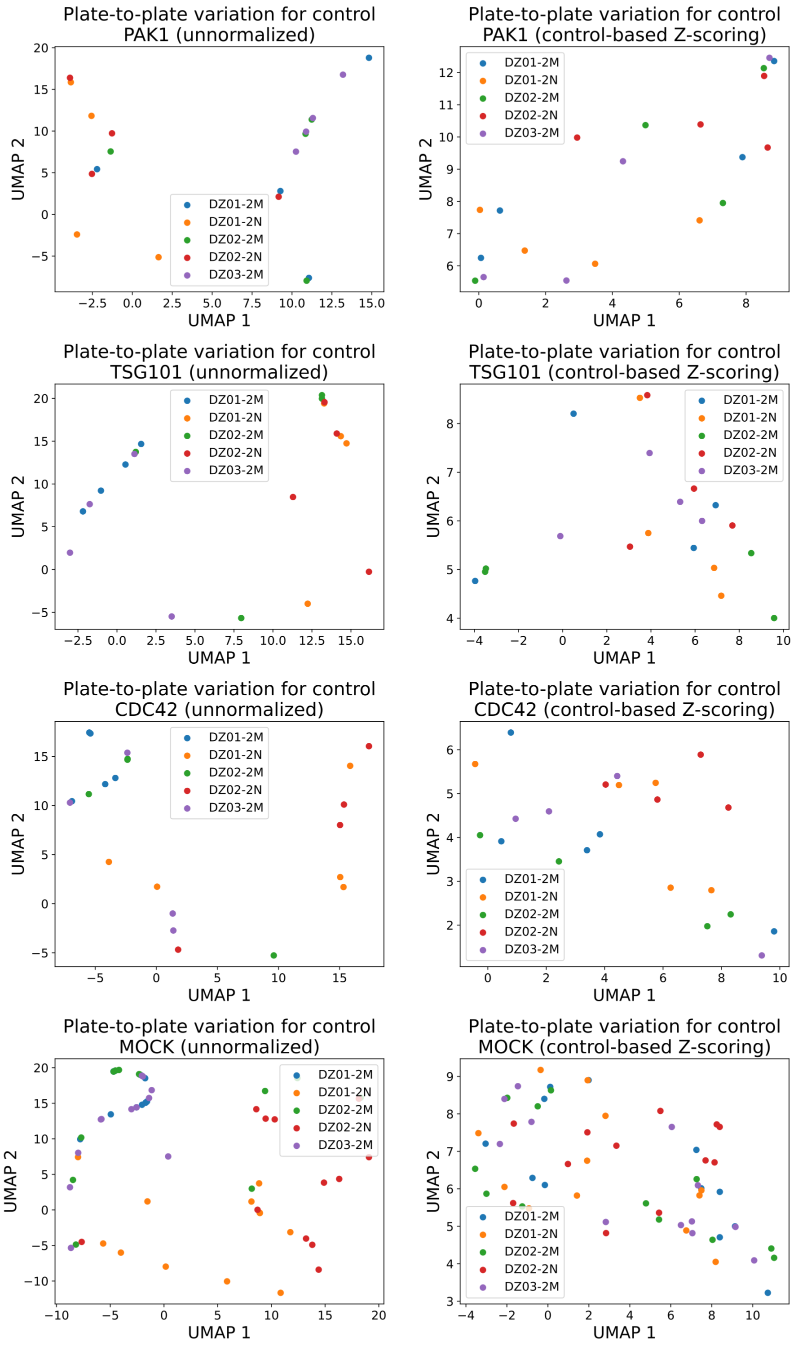


**Figure S1: UMAP visualisation of selected controls before and after control-based Z-scoring.**

### Identification of the most stable measurement type

To identify the most stable fluorescence intensity readout, defined as the readout exhibiting the lowest variability across plates, the coefficient of variation (CV; equation 2) was used:

$$\mathrm{CV} = \frac{\sigma}{\mu} \tag{2}$$

where $\sigma$ denotes the standard deviation and $\mu$ the mean. The CV expresses variability relative to the mean, making it scale-independent - a key consideration when comparing measurement types with different ranges.

Stability was assessed for each of the four fluorescence intensity readouts (whole-cell, nuclear, perinuclear and Voronoi-defined cellular regions) using a two-step procedure. First, for each control independently, the mean fluorescence intensity was calculated for every plate, and the corresponding cross-plate CV was determined. Second, the control-specific CV values were aggregated by calculating their median, yielding a single stability metric for each fluorescence intensity readout.

The controls included in the screen were ATP6V1A, SCRAMBLED, MOCK, PSMC3, KIF11, PSMA6, TSG101, GFP, RAC1, ARPC3, CDC42 and PAK1. Table S1 summarises the median CV values obtained for the different fluorescence intensity readouts.

**Table S1: Median CV values for the fluorescence intensity readouts**

| | Early viral gene expression (eGFP) | Late viral gene expression (mCherry) |
|---|---|---|
| Whole cell | 0.6487 | 0.9302 |
| Nuclear | 0.6947 | 0.6833 |
| Perinuclear | 0.6027 | 0.8410 |
| Voronoi-defined cellular regions | 0.6862 | 0.8543 |

Given the cytoplasmic replication cycle of VACV, fluorescence intensities derived from Voronoi-defined cellular regions were selected for quantification of both early and late viral gene expression.

#### Construction of the VACV-human PPI data set

Experimentally validated VACV-human protein-protein interactions (PPIs) were obtained from the Human-Virus Interaction Database (HVIDB). The initial dataset comprised 456 interactions across multiple vaccinia virus strains. Interactions were filtered to retain only those involving the Western Reserve (WR) strain. Following removal of three human proteins no longer present in UniProt, the final set of positive interactions comprised 412 PPIs.

Negative PPI instances were generated under two constraints. First, human proteins were required to localise exclusively to the nucleolus. Candidate proteins were retrieved from UniProt (organism: *Homo sapiens* [9606]; subcellular location: nucleolus [SL-0188]). As UniProt annotations may include multiple subcellular locations, entries were post-processed to retain only proteins exclusively annotated as nucleolar. One nucleolar protein (UniProt accession F4ZW62) present among the experimentally validated VACV interactors was excluded from the negative set.

Second, sequence dissimilarity between negative candidates and known interactors was enforced. Specifically, candidate proteins were required to exhibit less than 50% sequence identity to all human proteins involved in experimentally validated VACV PPIs. To this end, a multiple sequence alignment was performed using Clustal Omega (version 1.2.4; European Bioinformatics Institute web service) to compute a percent identity matrix.

To ensure compatibility with downstream structural analyses, all proteins included in the dataset were required to have a sequence length of at most 1,700 amino acids.

To minimise data leakage, the dataset was partitioned into training, validation and test sets using a combination of sequence-based clustering and interaction network structure. Human proteins eligible for negative sampling were clustered based on sequence identity, and clusters were assigned exclusively to one of the three subsets. In parallel, experimentally validated PPIs were represented as a bipartite graph, and connected components – comprising both viral and human proteins – were identified and assigned to a single subset to prevent overlap between splits.

Negative PPIs were generated independently within each subset by randomly pairing VACV WR proteins with the corresponding human nucleolar proteins. The resulting datasets were balanced to achieve a 1:1 ratio of positive to negative interactions. The final dataset comprised 686 PPIs in the training set, 68 PPIs in the validation set and 70 PPIs in the test set.

### Fine-tuning xCAPT5 with MLP classification head on the VACV PPI data set

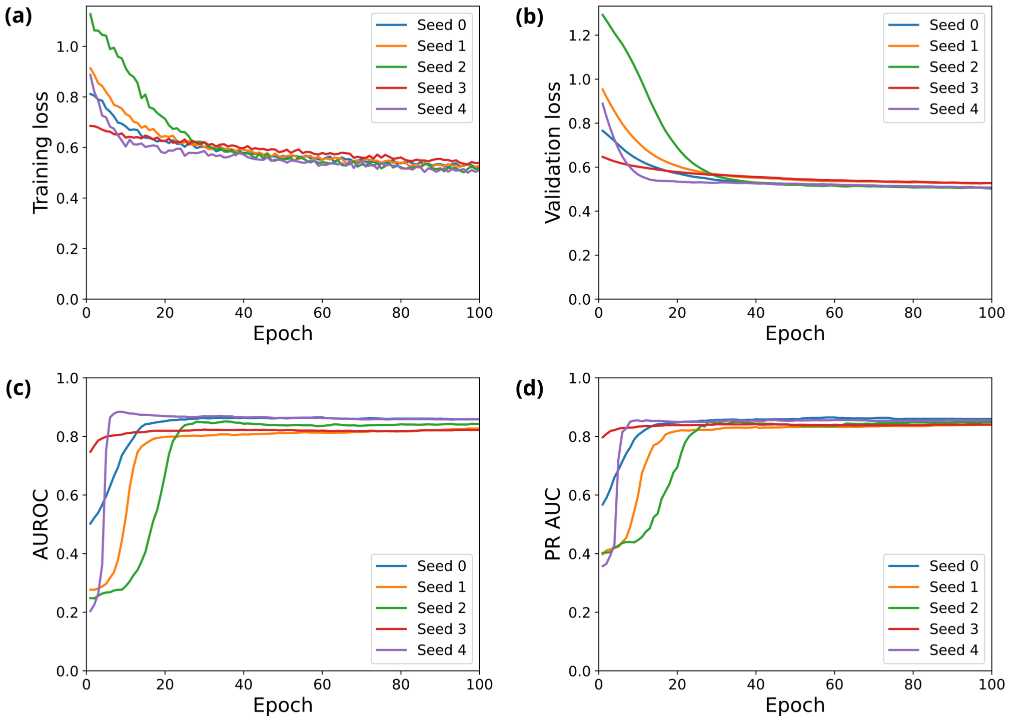


**Figure S2: Visualisation of training and validation metrics for the xCAPT5 Pan checkpoint.** (a) Training loss (b) Validation loss (c) Validation AUROC (d) Validation PR AUC

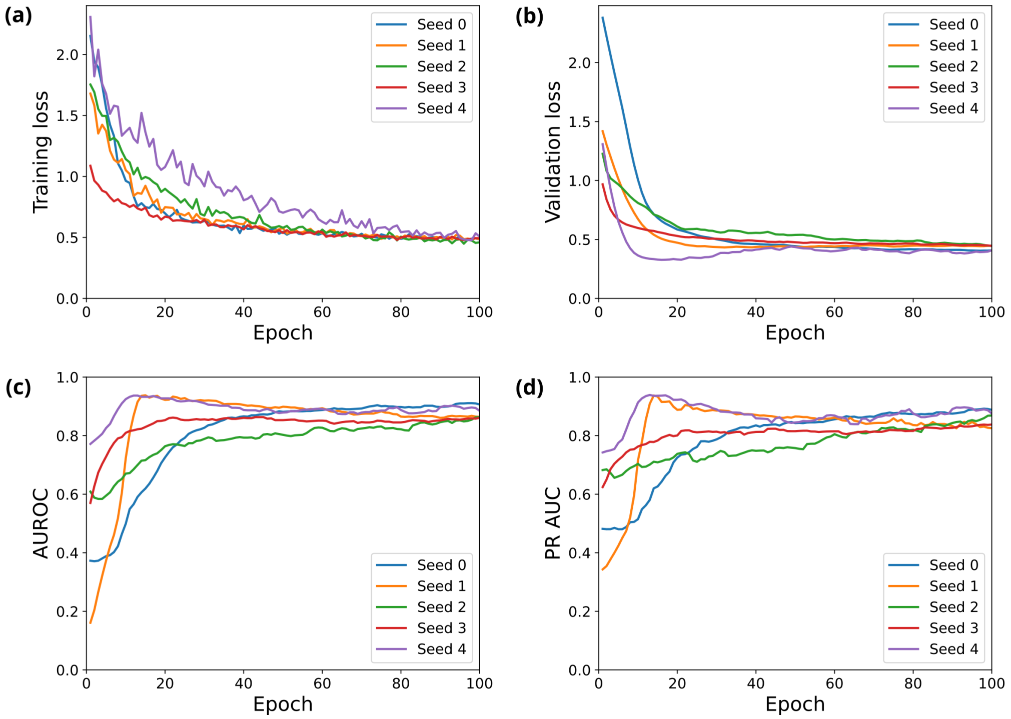


**Figure S3: Visualisation of training and validation metrics for the xCAPT5 Sled checkpoint.** (a) Training loss (b) Validation loss (c) Validation AUROC (d) Validation PR AUC

#### Detailed implementation of the construction of the PU learning data set

The framework requires genes to encode proteins. For this reason, the genes interrogated in the screen were filtered to retain only those associated with at least one encoded protein. This filtering step reduced the number of genes from 17.962 to 17.672.

Phenotype clustering was performed using k-means clustering on standardised phenotype feature vectors derived from the siRNA screen. Prior to clustering, phenotype vectors were Z-score normalised using standard scaling. The number of phenotype clusters was set to 10. Cluster assignments were subsequently propagated to the gene level.

To avoid information leakage, genes and proteins were represented as a bipartite graph in which genes were connected to all proteins encoded by the respective gene. Connected components of the graph were identified using breadth-first search (BFS) traversal and treated as atomic units during data set splitting. This ensured that neither genes nor proteins were shared across training, validation and test sets.

For each connected component, metadata including component size, phenotype cluster assignment, and host factor label were computed. Component-level splitting was then performed using a greedy balancing strategy. Components were sorted in descending order of component size. Initial split seeding ensured that each split contained at least one connected component. Remaining connected components were iteratively assigned to the split exhibiting the largest deficit relative to the predefined target split sizes. Target split proportions were set to 70% for training, 10% for validation and 20% for testing.

Because the downstream learning task operates at the gene level, all instances were collapsed to unique gene-level representations prior to splitting. Protein-level PPI feature vectors were subsequently aggregated to the gene level by max pooling of all protein-specific VACV interaction probability vectors associated with the respective gene.

To address class imbalance in the PU learning setting, a fixed positive-to-unlabelled (P:U) ratio of 1:5 was enforced independently within each split. Unlabelled genes were sampled in a phenotype-stratified manner according to phenotype cluster membership to preserve the distribution of phenotypic states across splits.

Phenotype distribution similarity between training, validation and test sets was quantified using the Jensen-Shannon divergence. The Jensen-Shannon divergence between the training and validation set and between the training and test set is 0.0806 and 0.0333, respectively, indicating a high degree of similarity between the individual phenotype distributions. Final splits were additionally validated to ensure one unique instance per gene, absence of gene overlap between splits and absence of protein overlap between splits.

All clustering, sampling and splitting procedures were performed using a fixed random seed to ensure reproducibility. Table S2 outlines the detailed composition of the PU learning data set.

**Table S2: Detailed composition of the PU learning data set.**

| Split | Positive instances | Unlabelled instances | Total |
|---|---|---|---|
| Training | 175 | 871 | 1.046 |
| Validation | 18 | 86 | 104 |
| Test | 50 | 244 | 294 |
| Total | 243 | 1.201 | 1.444 |

#### Architecture of the multimodal PU learning framework

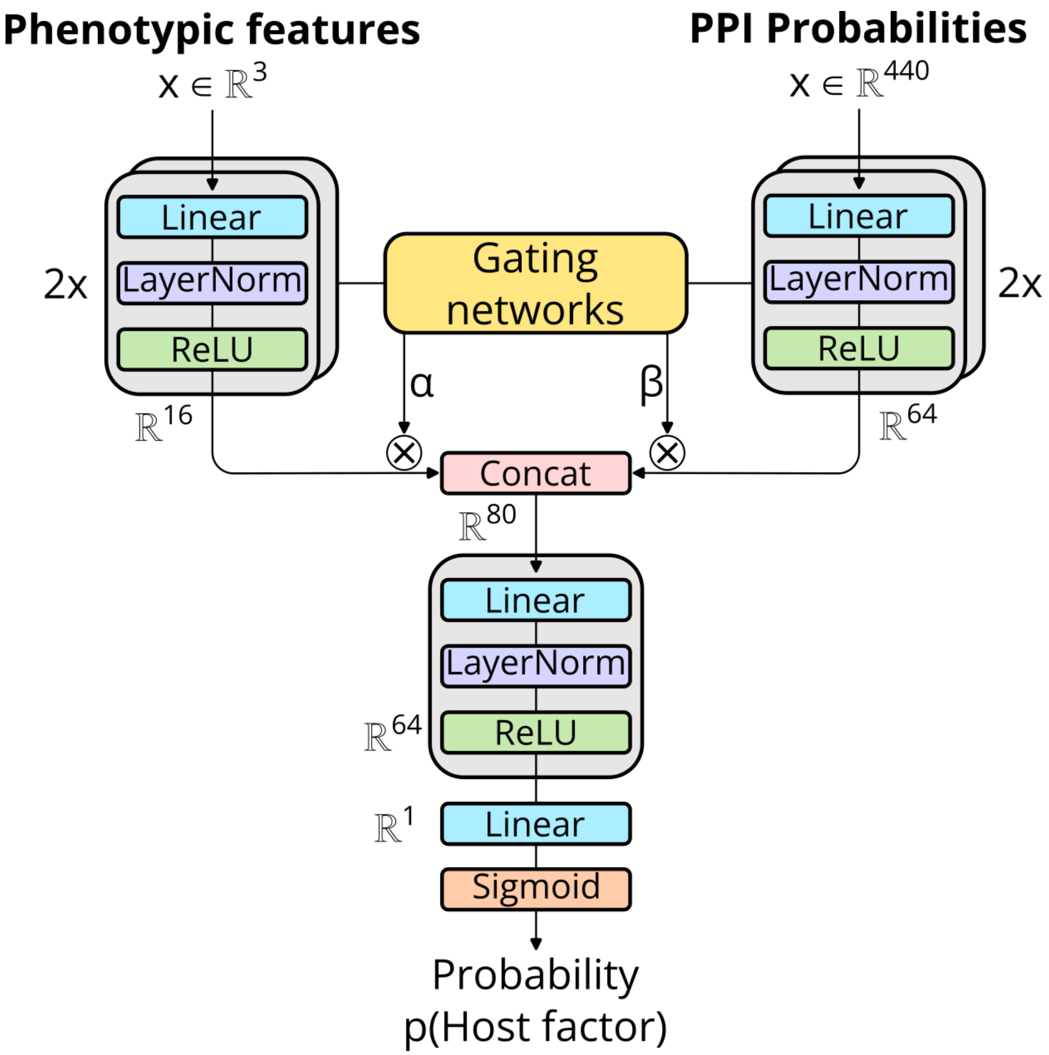


**Figure S4: Architecture of the multimodal PU learning framework.**

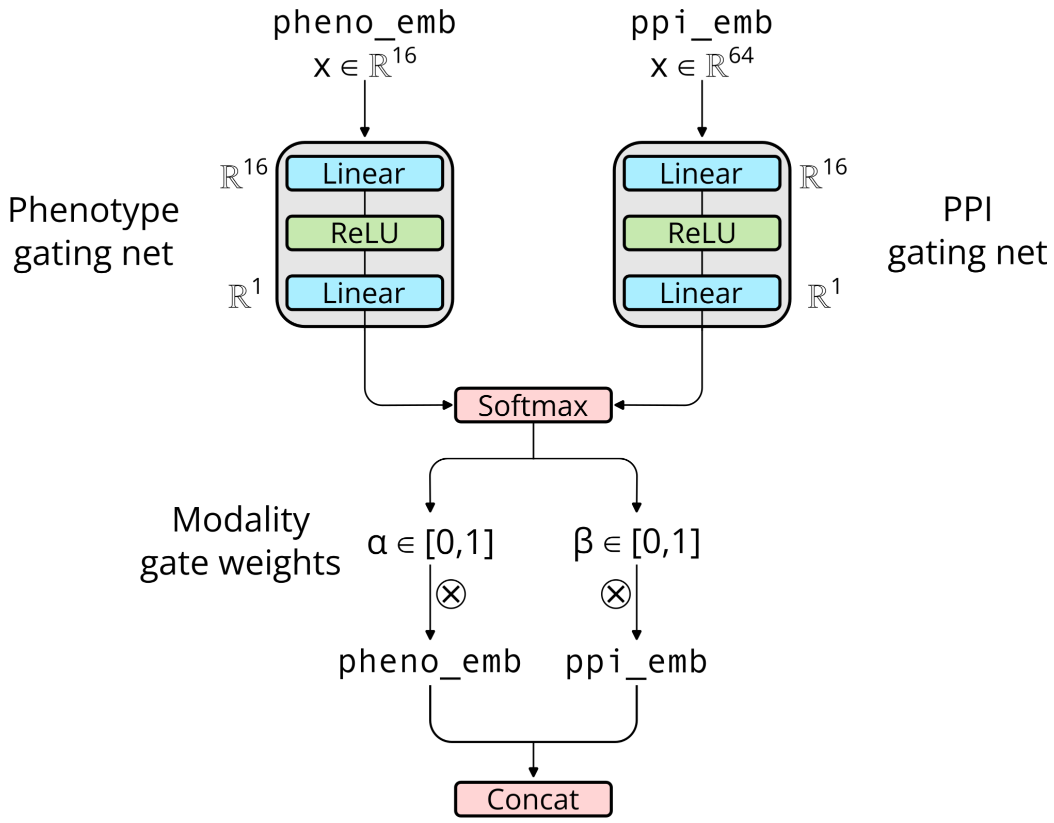


**Figure S5: Sample-specific modality-gating mechanism.**

### Training and evaluation of the PU learning framework

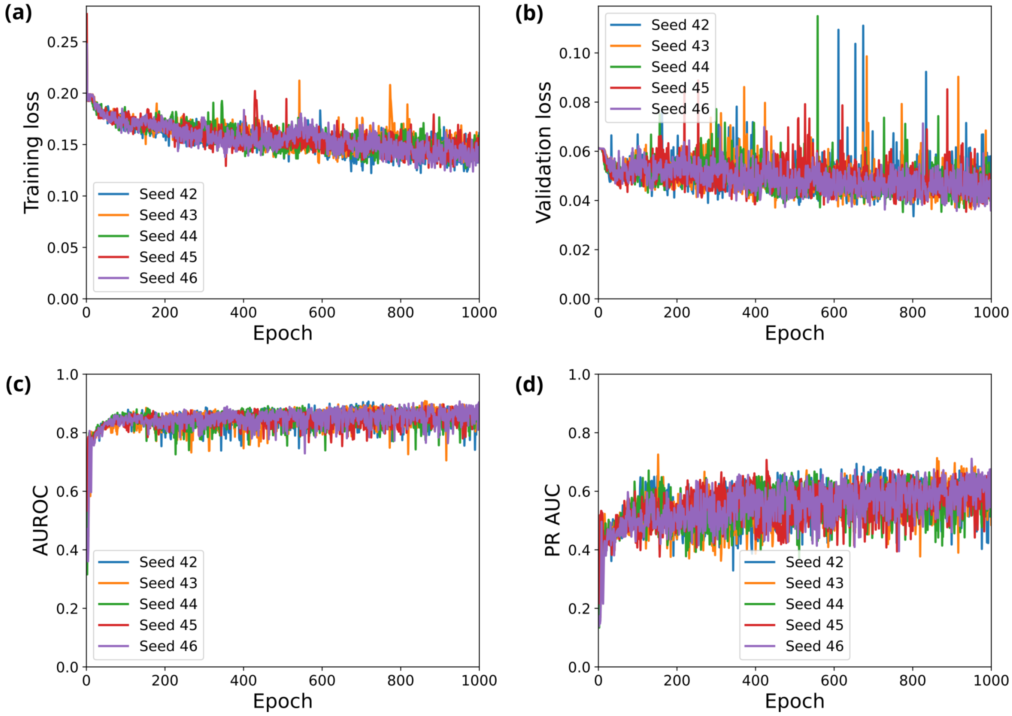


**Figure S6: Training and validation metrics for the PU learning architecture trained only on PPI features.** (a) Training loss (b) Validation loss (c) Validation AUROC (d) Validation PR AUC

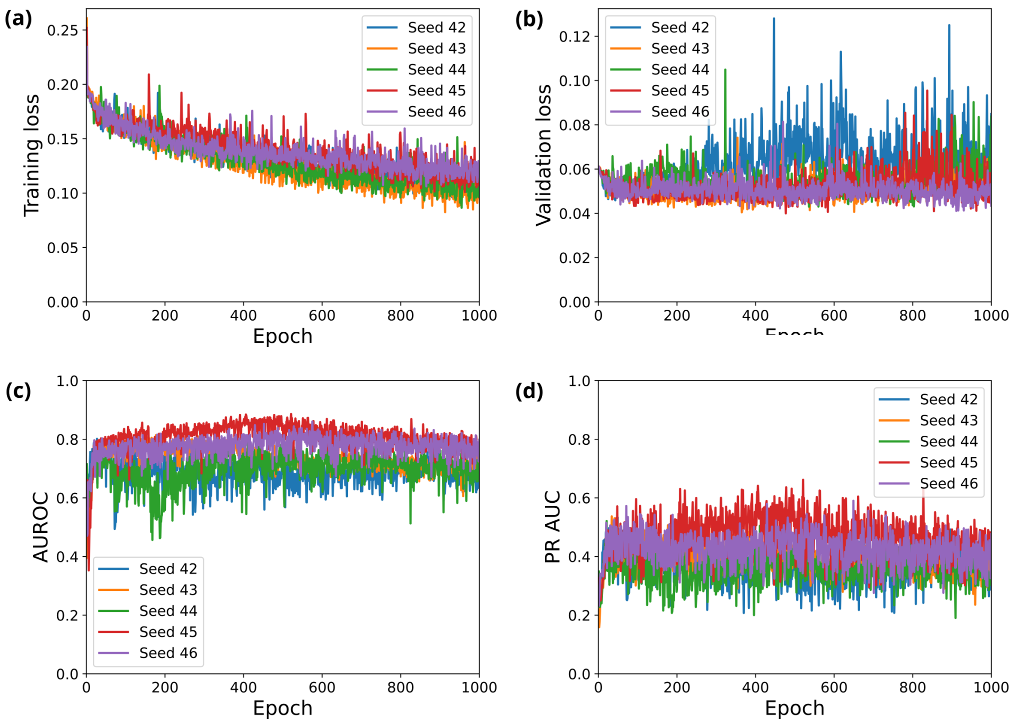


**Figure S7: Training and validation metrics for the PU learning architecture trained on both phenotypic and PPI features.** (a) Training loss (b) Validation loss (c) Validation AUROC (d) Validation PR AUC

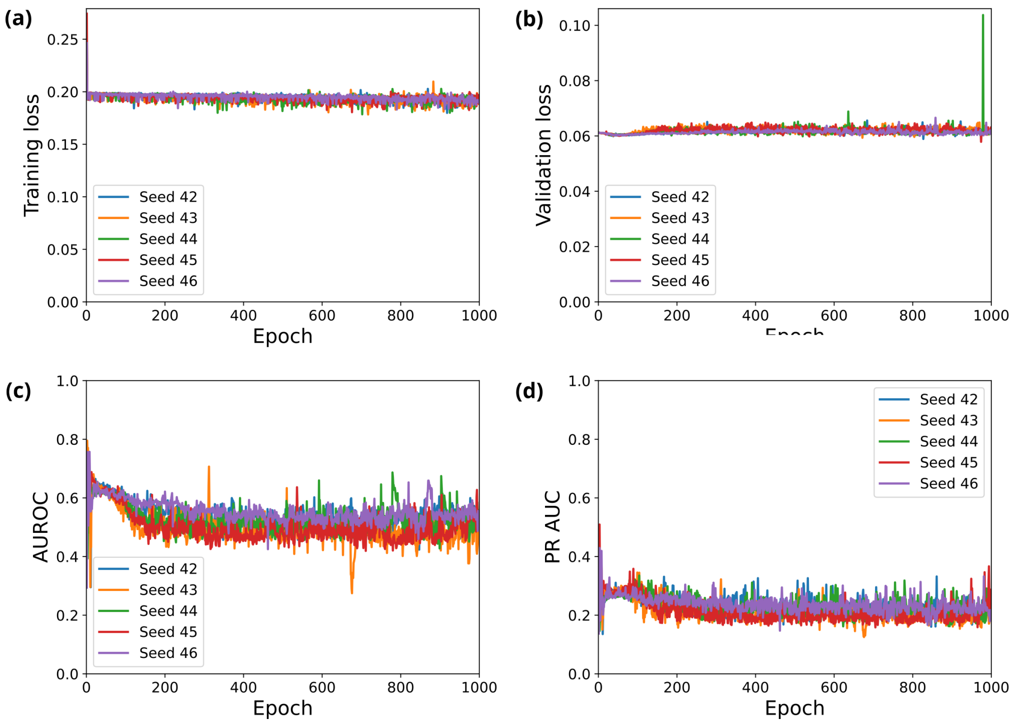


**Figure S8: Training and validation metrics for the PU learning architecture trained on row-permuted PPI features.** (a) Training loss (b) Validation loss (c) Validation AUROC (d) Validation PR AUC

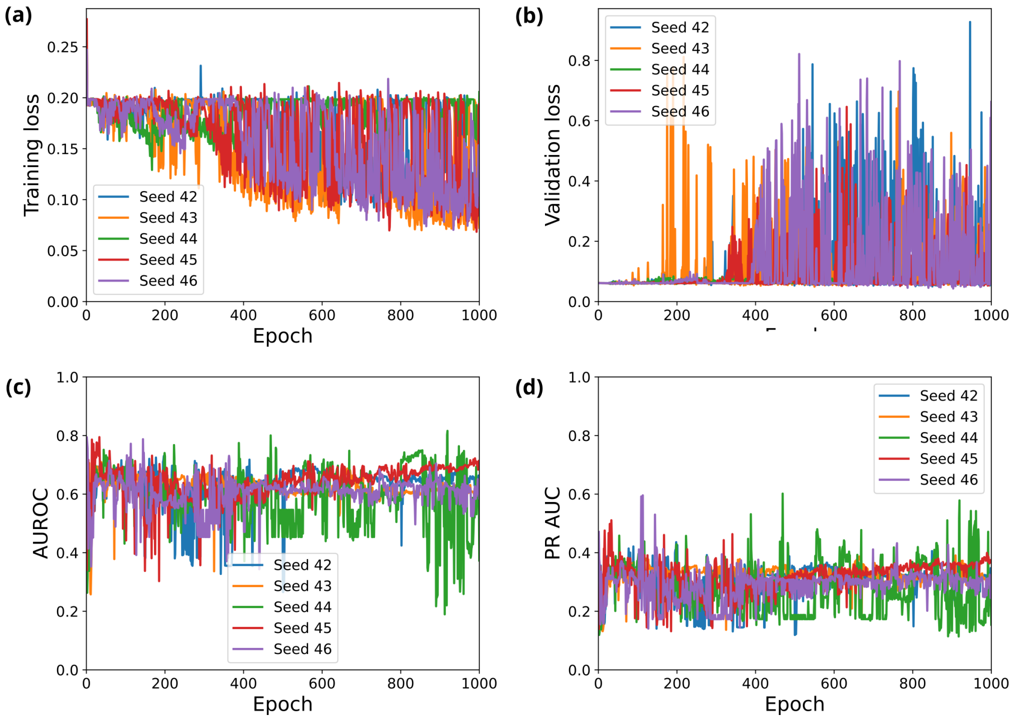


**Figure S9: Training and validation metrics for the PU learning architecture trained on column-permuted PPI features.** (a) Training loss (b) Validation loss (c) Validation AUROC (d) Validation PR AUC

**Table S3: Performance of the PU learning framework on the test set under different training settings.** Values represent mean ± s.d. across five random initialisations. Best performance in bold.

| | AUROC | PR AUC |
|---|---|---|
| Uncorrupted PPI features | **0.7732 ± 0.0065** | **0.4506 ± 0.0085** |
| Phenotypic + uncorrupted PPI features | 0.7174 ± 0.0201 | 0.4144 ± 0.0342 |
| Row-permuted PPI features | 0.4682 ± 0.0253 | 0.2000 ± 0.0321 |
| Column-permuted PPI features | 0.5049 ± 0.0122 | 0.2283 ± 0.0312 |

### Performance of different training configurations of the multimodal MLP on the PU test set

**Table S4: Top-k performance metrics for the PPI-only model (un-corrupted PPI features).** Values have been computed across five random initialisations.

| k | #positives _mean | #positives _std | recall @k_mean | recall @k_std | precision @k_mean | precision @k_std | enrichment @k_mean | enrichment @k_std |
|---|---|---|---|---|---|---|---|---|
| 10 | 6.0 | 0.6325 | 0.1200 | 0.0126 | 0.6000 | 0.0632 | 3.5280 | 0.3719 |
| 20 | 10.8 | 1.1662 | 0.2160 | 0.0233 | 0.5400 | 0.0583 | 3.1752 | 0.3429 |
| 30 | 15.6 | 0.8000 | 0.3120 | 0.0160 | 0.5200 | 0.0267 | 3.0576 | 0.1568 |
| 40 | 19.0 | 1.5492 | 0.3800 | 0.0310 | 0.4750 | 0.0387 | 2.7930 | 0.2277 |
| 50 | 23.0 | 1.6733 | 0.4600 | 0.0335 | 0.4600 | 0.0335 | 2.7048 | 0.1968 |
| 60 | 26.2 | 1.1662 | 0.5240 | 0.0233 | 0.4367 | 0.0194 | 2.5676 | 0.1143 |
| 70 | 28.8 | 0.7483 | 0.5760 | 0.0150 | 0.4114 | 0.0107 | 2.4192 | 0.0629 |
| 80 | 31.4 | 1.0198 | 0.6280 | 0.0204 | 0.3925 | 0.0127 | 2.3079 | 0.0750 |
| 90 | 34.6 | 0.8000 | 0.6920 | 0.0160 | 0.3844 | 0.0089 | 2.2605 | 0.0523 |
| 100 | 36.0 | 1.0954 | 0.7200 | 0.0219 | 0.3600 | 0.0110 | 2.1168 | 0.0644 |
| 110 | 37.2 | 1.1662 | 0.7440 | 0.0233 | 0.3382 | 0.0106 | 1.9885 | 0.0623 |
| 120 | 38.2 | 1.7205 | 0.7640 | 0.0344 | 0.3183 | 0.0143 | 1.8718 | 0.0843 |
| 130 | 39.2 | 0.7483 | 0.7840 | 0.0150 | 0.3015 | 0.0058 | 1.7730 | 0.0338 |
| 140 | 40.6 | 0.8000 | 0.8120 | 0.0160 | 0.2900 | 0.0057 | 1.7052 | 0.0336 |
| 150 | 41.6 | 0.4899 | 0.8320 | 0.0098 | 0.2773 | 0.0033 | 1.6307 | 0.0192 |
| 160 | 42.4 | 0.4899 | 0.8480 | 0.0098 | 0.2650 | 0.0031 | 1.5582 | 0.0180 |
| 170 | 42.8 | 0.4000 | 0.8560 | 0.0080 | 0.2518 | 0.0024 | 1.4804 | 0.0138 |
| 180 | 44.2 | 0.4000 | 0.8840 | 0.0080 | 0.2456 | 0.0022 | 1.4439 | 0.0131 |
| 190 | 44.6 | 0.4899 | 0.8920 | 0.0098 | 0.2347 | 0.0026 | 1.3803 | 0.0152 |
| 200 | 44.8 | 0.4000 | 0.8960 | 0.0080 | 0.2240 | 0.0020 | 1.3171 | 0.0118 |

**Table S5: Top-k performance metrics for the multimodal model (phenotypic + PPI features).** Values have been computed across five random initialisations.

| k | #positives _mean | #positives _std | recall @k_mean | recall @k_std | precision @k_mean | precision @k_std | enrichment @k_mean | enrichment @k_std |
|---|---|---|---|---|---|---|---|---|
| 10 | 7.0 | 0.6325 | 0.1400 | 0.0126 | 0.7000 | 0.0632 | 4.1160 | 0.3719 |
| 20 | 10.6 | 1.9596 | 0.2120 | 0.0392 | 0.5300 | 0.0980 | 3.1164 | 0.5761 |
| 30 | 12.4 | 2.7276 | 0.2480 | 0.0546 | 0.4133 | 0.0909 | 2.4304 | 0.5346 |
| 40 | 15.6 | 1.6248 | 0.3120 | 0.0325 | 0.3900 | 0.0406 | 2.2932 | 0.2388 |
| 50 | 18.0 | 2.1909 | 0.3600 | 0.0438 | 0.3600 | 0.0438 | 2.1168 | 0.2576 |
| 60 | 21.0 | 2.0976 | 0.4200 | 0.0420 | 0.3500 | 0.0350 | 2.0580 | 0.2056 |
| 70 | 23.2 | 2.1354 | 0.4640 | 0.0427 | 0.3314 | 0.0305 | 1.9488 | 0.1794 |
| 80 | 25.0 | 2.2804 | 0.5000 | 0.0456 | 0.3125 | 0.0285 | 1.8375 | 0.1676 |
| 90 | 27.6 | 2.9394 | 0.5520 | 0.0588 | 0.3067 | 0.0327 | 1.8032 | 0.1920 |
| 100 | 29.6 | 1.7436 | 0.5920 | 0.0349 | 0.2960 | 0.0174 | 1.7405 | 0.1025 |
| 110 | 32.0 | 1.8974 | 0.6400 | 0.0379 | 0.2909 | 0.0172 | 1.7105 | 0.1014 |
| 120 | 33.4 | 1.6248 | 0.6680 | 0.0325 | 0.2783 | 0.0135 | 1.6366 | 0.0796 |
| 130 | 34.6 | 1.2000 | 0.6920 | 0.0240 | 0.2662 | 0.0092 | 1.5650 | 0.0543 |
| 140 | 35.8 | 1.3266 | 0.7160 | 0.0265 | 0.2557 | 0.0095 | 1.5036 | 0.0557 |
| 150 | 37.2 | 0.7483 | 0.7440 | 0.0150 | 0.2480 | 0.0050 | 1.4582 | 0.0293 |
| 160 | 39.0 | 1.2649 | 0.7800 | 0.0253 | 0.2438 | 0.0079 | 1.4333 | 0.0465 |
| 170 | 41.6 | 1.0198 | 0.8320 | 0.0204 | 0.2447 | 0.0060 | 1.4389 | 0.0353 |
| 180 | 43.0 | 0.6325 | 0.8600 | 0.0126 | 0.2389 | 0.0035 | 1.4047 | 0.0207 |
| 190 | 44.0 | 0.8944 | 0.8800 | 0.0179 | 0.2316 | 0.0047 | 1.3617 | 0.0277 |
| 200 | 44.2 | 0.7483 | 0.8840 | 0.0150 | 0.2210 | 0.0037 | 1.2995 | 0.0220 |

**Table S6: Top-k performance metrics for the model trained on row-permuted PPI features.** Values have been computed across five random initialisations.

| k | #positives _mean | #positives _std | recall @k_mean | recall @k_std | precision @k_mean | precision @k_std | enrichment @k_mean | enrichment @k_std |
|---|---|---|---|---|---|---|---|---|
| 10 | 3.2 | 1.3266 | 0.0640 | 0.0265 | 0.3200 | 0.1327 | 1.8816 | 0.7801 |
| 20 | 5.6 | 1.2000 | 0.1120 | 0.0240 | 0.2800 | 0.0600 | 1.6464 | 0.3528 |
| 30 | 6.8 | 1.7205 | 0.1360 | 0.0344 | 0.2267 | 0.0573 | 1.3328 | 0.3372 |
| 40 | 8.0 | 2.0000 | 0.1600 | 0.0400 | 0.2000 | 0.0500 | 1.1760 | 0.2940 |
| 50 | 9.4 | 1.0198 | 0.1880 | 0.0204 | 0.1880 | 0.0204 | 1.1054 | 0.1199 |
| 60 | 10.4 | 1.0198 | 0.2080 | 0.0204 | 0.1733 | 0.0170 | 1.0192 | 0.0999 |
| 70 | 11.8 | 1.3266 | 0.2360 | 0.0265 | 0.1686 | 0.0190 | 0.9912 | 0.1114 |
| 80 | 13.0 | 1.4142 | 0.2600 | 0.0283 | 0.1625 | 0.0177 | 0.9555 | 0.1039 |
| 90 | 14.0 | 1.8974 | 0.2800 | 0.0379 | 0.1556 | 0.0211 | 0.9147 | 0.1240 |
| 100 | 15.8 | 2.3152 | 0.3160 | 0.0463 | 0.1580 | 0.0232 | 0.9290 | 0.1361 |
| 110 | 17.0 | 2.7568 | 0.3400 | 0.0551 | 0.1545 | 0.0251 | 0.9087 | 0.1474 |
| 120 | 18.6 | 2.9394 | 0.3720 | 0.0588 | 0.1550 | 0.0245 | 0.9114 | 0.1440 |
| 130 | 19.0 | 3.4641 | 0.3800 | 0.0693 | 0.1462 | 0.0266 | 0.8594 | 0.1567 |
| 140 | 20.2 | 2.9933 | 0.4040 | 0.0599 | 0.1443 | 0.0214 | 0.8484 | 0.1257 |
| 150 | 22.0 | 2.8284 | 0.4400 | 0.0566 | 0.1467 | 0.0189 | 0.8624 | 0.1109 |
| 160 | 24.2 | 1.9391 | 0.4840 | 0.0388 | 0.1513 | 0.0121 | 0.8894 | 0.0713 |
| 170 | 25.8 | 1.9391 | 0.5160 | 0.0388 | 0.1518 | 0.0114 | 0.8924 | 0.0671 |
| 180 | 27.6 | 1.4967 | 0.5520 | 0.0299 | 0.1533 | 0.0083 | 0.9016 | 0.0489 |
| 190 | 29.4 | 0.8000 | 0.5880 | 0.0160 | 0.1547 | 0.0042 | 0.9099 | 0.0248 |
| 200 | 32.8 | 0.4000 | 0.6560 | 0.0080 | 0.1640 | 0.0020 | 0.9643 | 0.0118 |

**Table S7: Top-k performance metrics for the model trained on column-permuted PPI features.** Values have been computed across five random initialisations.

| k | #positives _mean | #positives _std | recall @k_mean | recall @k_std | precision @k_mean | precision @k_std | enrichment @k_mean | enrichment @k_std |
|---|---|---|---|---|---|---|---|---|
| 10 | 3.2 | 1.1662 | 0.0640 | 0.0233 | 0.3200 | 0.1166 | 1.8816 | 0.6857 |
| 20 | 6.0 | 2.6077 | 0.1200 | 0.0522 | 0.3000 | 0.1304 | 1.7640 | 0.7667 |
| 30 | 8.4 | 3.7202 | 0.1680 | 0.0744 | 0.2800 | 0.1240 | 1.6464 | 0.7292 |
| 40 | 9.4 | 4.3174 | 0.1880 | 0.0863 | 0.2350 | 0.1079 | 1.3818 | 0.6347 |
| 50 | 11.2 | 5.1536 | 0.2240 | 0.1031 | 0.2240 | 0.1031 | 1.3171 | 0.6061 |
| 60 | 12.4 | 5.7827 | 0.2480 | 0.1157 | 0.2067 | 0.0964 | 1.2152 | 0.5667 |
| 70 | 13.0 | 6.0992 | 0.2600 | 0.1220 | 0.1857 | 0.0871 | 1.0920 | 0.5123 |
| 80 | 14.0 | 6.0992 | 0.2800 | 0.1220 | 0.1750 | 0.0762 | 1.0290 | 0.4483 |
| 90 | 14.6 | 6.3750 | 0.2920 | 0.1275 | 0.1622 | 0.0708 | 0.9539 | 0.4165 |
| 100 | 15.4 | 6.7409 | 0.3080 | 0.1348 | 0.1540 | 0.0674 | 0.9055 | 0.3964 |
| 110 | 15.6 | 6.8877 | 0.3120 | 0.1378 | 0.1418 | 0.0626 | 0.8339 | 0.3682 |
| 120 | 16.4 | 7.3919 | 0.3280 | 0.1478 | 0.1367 | 0.0616 | 0.8036 | 0.3622 |
| 130 | 17.8 | 8.0100 | 0.3560 | 0.1602 | 0.1369 | 0.0616 | 0.8051 | 0.3623 |
| 140 | 18.8 | 8.6116 | 0.3760 | 0.1722 | 0.1343 | 0.0615 | 0.7896 | 0.3617 |
| 150 | 19.4 | 8.3570 | 0.3880 | 0.1671 | 0.1293 | 0.0557 | 0.7605 | 0.3276 |
| 160 | 22.4 | 6.7409 | 0.4480 | 0.1348 | 0.1400 | 0.0421 | 0.8232 | 0.2477 |
| 170 | 24.8 | 5.4185 | 0.4960 | 0.1084 | 0.1459 | 0.0319 | 0.8578 | 0.1874 |
| 180 | 26.0 | 6.1968 | 0.5200 | 0.1239 | 0.1444 | 0.0344 | 0.8493 | 0.2024 |
| 190 | 28.4 | 4.5869 | 0.5680 | 0.0917 | 0.1495 | 0.0241 | 0.8789 | 0.1420 |
| 200 | 32.0 | 1.6733 | 0.6400 | 0.0335 | 0.1600 | 0.0084 | 0.9408 | 0.0492 |

### Confirmed VACV host factors (version 1)

**Table S8: Confirmed VACV host factors with literature references.**

| Confirmed VACV host factor | Reference(s) |
|---|---|
| ABL1 | [28] |
| ABL2 | [28] |
| ACTA1 | [9] |
| ACTA2 | [9] |
| ACTB | [9] |
| ACTC1 | [9] |
| ACTG1 | [9] |
| ACTG2 | [9] |
| ACTR2 | [2] |
| ACTR3 | [2] |
| AKT1 | [23] |
| ANXA2 | [27] |
| ARPC1A | [2] |
| ARPC1B | [2] |
| ARPC2 | [2] |
| ARPC3 | [2] |
| ARPC4 | [2] |
| ARPC5 | [2] |
| ARPC5L | [2] |
| ATP6AP1 | [60] |
| ATP6AP2 | [60] |
| ATP6V0A1 | [60] |
| ATP6V0A2 | [60, 62] |
| ATP6V0A4 | [60] |
| ATP6V0B | [60] |
| ATP6V0C | [60] |
| ATP6V0D1 | [60] |
| ATP6V0D2 | [60] |
| ATP6V0E1 | [60] |
| ATP6V0E2 | [60] |
| ATP6V1A | [60, 62] |

| | |
|---|---|
| ATP6V1B1 | [60] |
| ATP6V1B2 | [60] |
| ATP6V1C1 | [60] |
| ATP6V1C2 | [60] |
| ATP6V1D | [60] |
| ATP6V1E1 | [60] |
| ATP6V1E2 | [60] |
| ATP6V1F | [60] |
| ATP6V1G1 | [60] |
| ATP6V1G2 | [60] |
| ATP6V1G3 | [60] |
| ATP6V1H | [60] |
| ATR | [47] |
| AXL | [37] |
| B2M | [37] |
| BANF1 | [22] |
| CAPRIN1 | [26] |
| CARD9 | [52] |
| CCR5 | [48] |
| CDC42 | [28, 19] |
| CHEK1 | [47] |
| CHMP1A | [21] |
| CHMP3 | [21] |
| CHMP4C | [21] |
| CHMP6 | [21] |
| CLTA | [54] |
| CLTB | [54] |
| CLTC | [54] |
| COG1 | [35, 50] |
| COG2 | [35, 50] |
| COG3 | [35, 50] |
| COG4 | [35, 50] |
| COG5 | [35, 50] |
| COG6 | [35, 50] |
| COG7 | [35, 50] |
| COG8 | [35, 50] |

| | |
|---|---|
| COPB1 | [64, 30] |
| COPB2 | [64, 30] |
| COPE | [64] |
| CSNK2B | [1] |
| CUL3 | [41] |
| DDX3X | [16] |
| EGFR | [13] |
| EGR1 | [3] |
| EIF2AK2 | [45] |
| EIF4E | [12] |
| EIF4G1 | [63] |
| ELK1 | [3] |
| EXT1 | [35] |
| FHOD1 | [2, 28] |
| FLNB | [14] |
| G3BP1 | [26] |
| G3BP2 | [26] |
| GAS6 | [42] |
| GRB2 | [53, 4] |
| HSP90AB1 | [20] |
| HSPA8 | [24] |
| IFIT1 | [34] |
| IFIT2 | [34] |
| IFIT3 | [34] |
| IFITM3 | [29] |
| IFNA1 | [61] |
| IFNA10 | [61] |
| IFNA13 | [61] |
| IFNA14 | [61] |
| IFNA16 | [61] |
| IFNA17 | [61] |
| IFNA2 | [61] |
| IFNA21 | [61] |
| IFNA4 | [61] |
| IFNA5 | [61] |
| IFNA6 | [61] |

| | |
|---|---|
| IFNA7 | [61] |
| IFNA8 | [61] |
| IFNB1 | [61] |
| IFNG | [61] |
| IL18 | [49, 15, 7] |
| IL1A | [7] |
| IL1B | [15] |
| INTS7 | [47] |
| ISG15 | [6] |
| ITGB1 | [37, 23] |
| ITSN1 | [19] |
| KDELR1 | [64] |
| KDELR2 | [64] |
| KDELR3 | [64] |
| KPNA2 | [46, 65] |
| LAMA1 | [41] |
| LAMA2 | [41] |
| MAP2K1 | [11] |
| MAP2K2 | [11] |
| MAP2K3 | [11] |
| MAP2K4 | [11] |
| MAP2K5 | [11] |
| MAP2K6 | [11] |
| MAP2K7 | [11] |
| MAPK1 | [11] |
| MAPK3 | [11] |
| MTOR | [39] |
| MYD88 | [7, 58] |
| MYO9A | [28] |
| NCK1 | [53] |
| NCK2 | [53] |
| NLRP1 | [15] |
| PAK1 | [40] |
| PCNA | [47] |
| PDCD6IP | [21] |
| PFN1 | [2, 31] |

| | |
|---|---|
| PIK3CA | [23, 57] |
| PIK3CB | [23] |
| PIK3R1 | [38] |
| PIK3R2 | [38] |
| PIK3R4 | [51] |
| PIKFYVE | [51] |
| PRKAA1 | [43] |
| PRKAA2 | [43, 5] |
| PRKAB1 | [43, 5] |
| PRKAB2 | [43, 5] |
| PRKACA | [11] |
| PRKACB | [11] |
| PRKACG | [11] |
| PRKAG1 | [43] |
| PRKAG2 | [43] |
| PRKAG3 | [43] |
| PRKAR1A | [11] |
| PRKAR1B | [11] |
| PRKAR2A | [11] |
| PRKAR2B | [11] |
| PRKDC | [55] |
| PSMA1 | [41] |
| PSMA2 | [41] |
| PSMA3 | [41] |
| PSMA4 | [41] |
| PSMA5 | [41] |
| PSMA6 | [41] |
| PSMA7 | [41] |
| PSMA8 | [41] |
| PSMB1 | [41] |
| PSMB10 | [41] |
| PSMB2 | [41] |
| PSMB3 | [41] |
| PSMB4 | [41] |
| PSMB5 | [41] |
| PSMB6 | [41] |

| | |
|---|---|
| PSMB7 | [41] |
| PSMB8 | [41] |
| PSMB9 | [41] |
| PSMC1 | [41] |
| PSMC2 | [41] |
| PSMC3 | [41] |
| PSMC4 | [41] |
| PSMC5 | [41] |
| PSMC6 | [41] |
| PSMD1 | [41] |
| PSMD10 | [41] |
| PSMD11 | [41] |
| PSMD12 | [41] |
| PSMD13 | [41] |
| PSMD14 | [41] |
| PSMD2 | [41] |
| PSMD3 | [41] |
| PSMD4 | [41] |
| PSMD5 | [41] |
| PSMD6 | [41] |
| PSMD7 | [41] |
| PSMD8 | [41] |
| PSMD9 | [41] |
| RAB1A | [25] |
| RAB34 | [51] |
| RAB5C | [51] |
| RAB7A | [51] |
| RABGEF1 | [51] |
| RAC1 | [2] |
| RAD50 | [52] |
| RBX1 | [41] |
| RHNO1 | [47] |
| RHOA | [28, 10] |
| RIPK3 | [8] |
| RPA2 | [47] |
| RPS6KA3 | [3] |

| | |
|---|---|
| SAMD9 | [33, 56] |
| SNX3 | [51] |
| SRC | [28] |
| STAT1 | [36] |
| STAT2 | [59] |
| TFAP2A | [54] |
| TFAP2B | [54] |
| TFAP2C | [54] |
| TFAP2D | [54] |
| TFAP2E | [54] |
| TICAM1 | [58] |
| TICAM2 | [58] |
| TM9SF2 | [35] |
| TMED10 | [35] |
| TNF | [61] |
| TOPBP1 | [47] |
| TSG101 | [21] |
| UBA1 | [41] |
| UBE2D1 | [44] |
| VEGFA | [17] |
| VIM | [16] |
| VIPAS39 | [30] |
| VPS26A | [30] |
| VPS26B | [30] |
| VPS33B | [30] |
| VPS35 | [30] |
| VPS4B | [21] |
| VRK1 | [32] |
| WASHC2C | [18] |
| WASL | [2] |
| WDR6 | [56] |
| XRCC5 | [55] |
| XRCC6 | [55] |

### Mapping of Reactome pathway terms to ten broad biological categories

**Table S9: Assignment of Reactome pathway terms to ten broad biological categories.**

| Reactome pathway term | Biological category |
|---|---|
| SMAD2/SMAD3:SMAD4 heterotrimer regulates transcription | Cell signaling |
| Eukaryotic Translation Initiation | Protein & RNA homeostasis |
| Inhibition of replication initiation of damaged DNA by RB1/E2F1 | DNA replication & repair |
| Alpha-protein kinase 1 signaling pathway | Cell signaling |
| Aggrephagy | Autophagy |
| p53-Dependent G1 DNA Damage Response | DNA replication & repair |
| Degradation of CDH1 | Cell cycle & mitosis |
| Cellular response to chemical stress | Other |
| Intracellular signaling by second messengers | Cell signaling |
| Defective homologous recombination repair (HRR) due to PALB2 loss of function | DNA replication & repair |
| Major pathway of rRNA processing in the nucleolus and cytosol | Protein & RNA homeostasis |
| RNA Polymerase II Transcription Initiation And Promoter Clearance | Protein & RNA homeostasis |
| G1/S Transition | Cell cycle & mitosis |
| Signaling by ERBB2 | Cell signaling |
| Mismatch repair (MMR) directed by MSH2:MSH6 (MutSalpha) | DNA replication & repair |
| Interactions of Rev with host cellular proteins | Immune response & host–pathogen interactions |

| | |
|---|---|
| HDR through Homologous Recombination (HRR) | DNA replication & repair |
| DNA replication initiation | DNA replication & repair |
| Gene expression (Transcription) | Protein & RNA homeostasis |
| SCF-beta-TrCP mediated degradation of Emi1 | Cell cycle & mitosis |
| Regulation of PTEN stability and activity | Cell signaling |
| L13a-mediated translational silencing of Ceruloplasmin expression | Protein & RNA homeostasis |
| Cell Cycle, Mitotic | Cell cycle & mitosis |
| Metabolism | Other |
| RNA Polymerase II Pre-transcription Events | Protein & RNA homeostasis |
| Calnexin/calreticulin cycle | Protein & RNA homeostasis |
| G0 and Early G1 | Cell cycle & mitosis |
| PKR-mediated signaling | Immune response & host–pathogen interactions |
| Global Genome Nucleotide Excision Repair (GG-NER) | DNA replication & repair |
| Formation of the HIV-1 Early Elongation Complex | Immune response & host–pathogen interactions |
| Pausing and recovery of HIV elongation | Immune response & host–pathogen interactions |
| SUMOylation of DNA replication proteins | DNA replication & repair |
| Cytosolic tRNA aminoacylation | Protein & RNA homeostasis |

| | |
|---|---|
| Ribosomal scanning and start codon recognition | Protein & RNA homeostasis |
| IRAK1 recruits IKK complex upon TLR7/8 or 9 stimulation | Immune response & host–pathogen interactions |
| Intra-Golgi and retrograde Golgi-to-ER traffic | Membrane trafficking & vesicular transport |
| mTORC1-mediated signalling | Cell signaling |
| Deubiquitination | Ubiquitin-mediated protein degradation |
| Cargo trafficking to the periciliary membrane | Membrane trafficking & vesicular transport |
| Abortive elongation of HIV-1 transcript in the absence of Tat | Immune response & host–pathogen interactions |
| Response of EIF2AK4 (GCN2) to amino acid deficiency | Cell signaling |
| Cell-cell junction organization | Cytoskeleton & Rho GTPase signaling |
| Transport of Mature mRNA Derived from an Intronless Transcript | Protein & RNA homeostasis |
| M-decay: degradation of maternal mRNAs by maternally stored factors | Protein & RNA homeostasis |
| PCP/CE pathway | Cell signaling |
| Mitotic Prophase | Cell cycle & mitosis |
| VLDLR internalisation and degradation | Membrane trafficking & vesicular transport |
| Regulation of BACH1 activity | Cell signaling |
| Autodegradation of the E3 ubiquitin ligase COP1 | Ubiquitin-mediated protein degradation |

| | |
|---|---|
| RNA Pol II CTD phosphorylation and interaction with CE | Protein & RNA homeostasis |
| Maturation of DENV proteins | Immune response & host–pathogen interactions |
| CHD chromatin remodelers | Protein & RNA homeostasis |
| mRNA 3'-end processing | Protein & RNA homeostasis |
| E2F mediated regulation of DNA replication | DNA replication & repair |
| Transcriptional regulation by small RNAs | Protein & RNA homeostasis |
| Chromatin organization | Protein & RNA homeostasis |
| Response of Mtb to phagocytosis | Immune response & host–pathogen interactions |
| Ubiquitin-Mediated Degradation of Phosphorylated Cdc25A | Cell cycle & mitosis |
| Centrosome maturation | Cell cycle & mitosis |
| Hedgehog 'off' state | Cell signaling |
| HIV Transcription Initiation | Immune response & host–pathogen interactions |
| Formation of HIV elongation complex in the absence of HIV Tat | Immune response & host–pathogen interactions |
| Regulation of TP53 Activity | Cell signaling |
| Removal of the Flap Intermediate from the C-strand | DNA replication & repair |
| Signaling by WNT | Cell signaling |
| Signaling by NOTCH | Cell signaling |

| | |
|---|---|
| Rev-mediated nuclear export of HIV RNA | Immune response & host–pathogen interactions |
| MAPK6/MAPK4 signaling | Cell signaling |
| Regulation of CDH1 Function | Cell cycle & mitosis |
| Nuclear events mediated by NFE2L2 | Cell signaling |
| Defective HDR through Homologous Recombination Repair (HRR) due to PALB2 loss of BRCA1 binding function | DNA replication & repair |
| Hh mutants are degraded by ERAD | Ubiquitin-mediated protein degradation |
| Downstream signaling events of B Cell Receptor (BCR) | Immune response & host–pathogen interactions |
| mRNA Splicing | Protein & RNA homeostasis |
| Cell-Cell communication | Other |
| Loss of proteins required for interphase microtubule organization from the centrosome | Cytoskeleton & Rho GTPase signaling |
| DNA Replication | DNA replication & repair |
| Josephin domain DUBs | Ubiquitin-mediated protein degradation |
| Signaling by ROBO receptors | Cell signaling |
| Autodegradation of Cdh1 by Cdh1:APC/C | Cell cycle & mitosis |
| rRNA processing in the nucleus and cytosol | Protein & RNA homeostasis |
| Termination of translesion DNA synthesis | DNA replication & repair |
| Impaired BRCA2 binding to RAD51 | DNA replication & repair |

| | |
|---|---|
| PD-L1(CD274) glycosylation and translocation to plasma membrane | Immune response & host–pathogen interactions |
| DNA Damage Recognition in GG-NER | DNA replication & repair |
| APC/C:Cdc20 mediated degradation of Securin | Cell cycle & mitosis |
| Synthesis of active ubiquitin: roles of E1 and E2 enzymes | Ubiquitin-mediated protein degradation |
| Developmental Biology | Other |
| Transport of Mature mRNAs Derived from Intronless Transcripts | Protein & RNA homeostasis |
| Golgi-to-ER retrograde transport | Membrane trafficking & vesicular transport |
| Regulation of TBK1, IKK$\epsilon$-mediated activation of IRF3, IRF7 upon TLR3 ligation | Immune response & host–pathogen interactions |
| EPHB-mediated forward signaling | Cell signaling |
| HIV Transcription Elongation | Immune response & host–pathogen interactions |
| Clathrin-mediated endocytosis | Membrane trafficking & vesicular transport |
| Regulation of activated PAK-2p34 by proteasome mediated degradation | Ubiquitin-mediated protein degradation |
| Immune System | Immune response & host–pathogen interactions |
| SARS-CoV-2 Infection | Immune response & host–pathogen interactions |

| | |
|---|---|
| TNFR2 non-canonical NF-kB pathway | Immune response & host–pathogen interactions |
| HuR (ELAVL1) binds and stabilizes mRNA | Protein & RNA homeostasis |
| Defective HDR through Homologous Recombination Repair (HRR) due to PALB2 loss of BRCA2/RAD51/RAD51C binding function | DNA replication & repair |
| Nonsense-Mediated Decay (NMD) | Protein & RNA homeostasis |
| Signaling by NOTCH4 | Cell signaling |
| Resolution of Abasic Sites (AP sites) | DNA replication & repair |
| mRNA Splicing - Major Pathway | Protein & RNA homeostasis |
| Inhibition of DNA recombination at telomere | DNA replication & repair |
| Cellular response to starvation | Autophagy |
| Telomere C-strand (Lagging Strand) Synthesis | DNA replication & repair |
| ER Quality Control Compartment (ERQC) | Protein & RNA homeostasis |
| G2/M Checkpoints | Cell cycle & mitosis |
| Formation of a pool of free 40S subunits | Protein & RNA homeostasis |
| Transport of Mature mRNA derived from an Intron-Containing Transcript | Protein & RNA homeostasis |
| Iron uptake and transport | Other |
| RUNX1 regulates transcription of genes involved in differentiation of HSCs | Cell signaling |
| Amplification of signal from unattached kinetochores via a MAD2 inhibitory signal | Cell cycle & mitosis |
| Nonsense Mediated Decay (NMD) independent of the Exon Junction Complex (EJC) | Protein & RNA homeostasis |

| | |
|---|---|
| Processing of DNA double-strand break ends | DNA replication & repair |
| NEP/NS2 Interacts with the Cellular Export Machinery | Immune response & host–pathogen interactions |
| Cyclin A:Cdk2-associated events at S phase entry | Cell cycle & mitosis |
| Asymmetric localization of PCP proteins | Cell signaling |
| Metabolism of proteins | Protein & RNA homeostasis |
| Mitophagy | Autophagy |
| Regulation of Expression and Function of Type I Classical Cadherins | Cytoskeleton & Rho GTPase signaling |
| Mitotic Spindle Checkpoint | Cell cycle & mitosis |
| Degradation of GLI2 by the proteasome | Cell signaling |
| G1/S-Specific Transcription | Cell cycle & mitosis |
| Generic Transcription Pathway | Protein & RNA homeostasis |
| Mitotic Prometaphase | Cell cycle & mitosis |
| NOTCH1 Intracellular Domain Regulates Transcription | Cell signaling |
| Constitutive Signaling by Ligand-Responsive EGFR Cancer Variants | Cell signaling |
| Infectious disease | Immune response & host–pathogen interactions |
| Deadenylation of mRNA | Protein & RNA homeostasis |
| Diseases of signal transduction by growth factor receptors and second messengers | Cell signaling |
| Resolution of AP sites via the multiple-nucleotide patch replacement pathway | DNA replication & repair |

| | |
|---|---|
| Regulation of TNFR1 signaling | Immune response & host–pathogen interactions |
| Fanconi Anemia Pathway | DNA replication & repair |
| Post-translational protein modification | Protein & RNA homeostasis |
| Transcription-Coupled Nucleotide Excision Repair (TC-NER) | DNA replication & repair |
| HIV elongation arrest and recovery | Immune response & host–pathogen interactions |
| tRNA processing | Protein & RNA homeostasis |
| Ribosome Quality Control (RQC) complex extracts and degrades nascent peptide | Protein & RNA homeostasis |
| Recruitment of mitotic centrosome proteins and complexes | Cell cycle & mitosis |
| SARS-CoV-1 modulates host translation machinery | Immune response & host–pathogen interactions |
| Infection with Mycobacterium tuberculosis | Immune response & host–pathogen interactions |
| Regulation of CDH1 Expression and Function | Cell cycle & mitosis |
| Vpu mediated degradation of CD4 | Immune response & host–pathogen interactions |
| Membrane Trafficking | Membrane trafficking & vesicular transport |
| Nonsense Mediated Decay (NMD) enhanced by the Exon Junction Complex (EJC) | Protein & RNA homeostasis |

| | |
|---|---|
| Downstream TCR signaling | Immune response & host–pathogen interactions |
| AURKA Activation by TPX2 | Cell cycle & mitosis |
| Pausing and recovery of Tat-mediated HIV elongation | Immune response & host–pathogen interactions |
| RNA Polymerase II Transcription | Protein & RNA homeostasis |
| Formation of the ternary complex, and subsequently, the 43S complex | Protein & RNA homeostasis |
| Formation of HIV-1 elongation complex containing HIV-1 Tat | Immune response & host–pathogen interactions |
| IRAK2 mediated activation of TAK1 complex | Immune response & host–pathogen interactions |
| TICAM1,TRAF6-dependent induction of TAK1 complex | Immune response & host–pathogen interactions |
| Regulation of TP53 Degradation | Cell signaling |
| Lagging Strand Synthesis | DNA replication & repair |
| Signaling by FGFR2 IIIa TM | Cell signaling |
| Signaling by Ligand-Responsive EGFR Variants in Cancer | Cell signaling |
| Hedgehog ligand biogenesis | Cell signaling |
| Evasion by RSV of host interferon responses | Immune response & host–pathogen interactions |
| PIP3 activates AKT signaling | Cell signaling |
| HSP90 chaperone cycle for steroid hormone receptors (SHR) in the presence of ligand | Protein & RNA homeostasis |
| Transcriptional regulation by RUNX3 | Cell signaling |

| | |
|---|---|
| Viral mRNA Translation | Immune response & host–pathogen interactions |
| Signaling by Hedgehog | Cell signaling |
| Transcriptional regulation by RUNX2 | Cell signaling |
| Transcriptional and post-translational regulation of MITF-M expression and activity | Cell signaling |
| Cooperation of Prefoldin and TriC/CCT in actin and tubulin folding | Cytoskeleton & Rho GTPase signaling |
| RAF/MAP kinase cascade | Cell signaling |
| Translesion synthesis by Y family DNA polymerases bypasses lesions on DNA template | DNA replication & repair |
| Defective homologous recombination repair (HRR) due to BRCA1 loss of function | DNA replication & repair |
| Regulation of mitotic cell cycle | Cell cycle & mitosis |
| Activation of STAT3 by cadherin engagement | Cell signaling |
| mRNA Splicing - Minor Pathway | Protein & RNA homeostasis |
| Axon guidance | Cytoskeleton & Rho GTPase signaling |
| AUF1 (hnRNP D0) binds and destabilizes mRNA | Protein & RNA homeostasis |
| Chaperonin-mediated protein folding | Protein & RNA homeostasis |
| TGF-beta receptor signaling activates SMADs | Cell signaling |
| Formation of Incision Complex in GG-NER | DNA replication & repair |
| Degradation of CRY and PER proteins | Ubiquitin-mediated protein degradation |
| Base Excision Repair | DNA replication & repair |

| | |
|---|---|
| Extension of Telomeres | DNA replication & repair |
| COPI-independent Golgi-to-ER retrograde traffic | Membrane trafficking & vesicular transport |
| Host Interactions of HIV factors | Immune response & host–pathogen interactions |
| Programmed Cell Death | Other |
| Transcriptional activity of SMAD2/SMAD3:SMAD4 heterotrimer | Cell signaling |
| Proteasome assembly | Ubiquitin-mediated protein degradation |
| APC/C:Cdc20 mediated degradation of Cyclin B | Cell cycle & mitosis |
| APC-Cdc20 mediated degradation of Nek2A | Cell cycle & mitosis |
| Signaling by the B Cell Receptor (BCR) | Immune response & host–pathogen interactions |
| Transferrin endocytosis and recycling | Membrane trafficking & vesicular transport |
| Amplification of signal from the kinetochores | Cell cycle & mitosis |
| Signaling by CSF3 (G-CSF) | Immune response & host–pathogen interactions |
| CHD1 and CHD2 subfamily | Protein & RNA homeostasis |
| RNA Polymerase II HIV Promoter Escape | Immune response & host–pathogen interactions |
| Mismatch repair (MMR) directed by MSH2:MSH3 (MutSbeta) | DNA replication & repair |

| | |
|---|---|
| RNA Polymerase II Transcription Initiation | Protein & RNA homeostasis |
| APC/C-mediated degradation of cell cycle proteins | Cell cycle & mitosis |
| Regulation of Homotypic Cell-Cell Adhesion | Cytoskeleton & Rho GTPase signaling |
| Tat-mediated elongation of the HIV-1 transcript | Immune response & host–pathogen interactions |
| Signal Transduction | Cell signaling |
| Asparagine N-linked glycosylation | Protein & RNA homeostasis |
| ZNF598 and the Ribosome-associated Quality Trigger (RQT) complex dissociate a ribosome stalled on a no-go mRNA | Protein & RNA homeostasis |
| JNK (c-Jun kinases) phosphorylation and activation mediated by activated human TAK1 | Cell signaling |
| snRNP Assembly | Protein & RNA homeostasis |
| Loss of Nlp from mitotic centrosomes | Cell cycle & mitosis |
| Beta-catenin independent WNT signaling | Cell signaling |
| Dengue Virus Infection | Immune response & host–pathogen interactions |
| Metalloprotease DUBs | Ubiquitin-mediated protein degradation |
| Pexophagy | Autophagy |
| BBSome-mediated cargo-targeting to cilium | Membrane trafficking & vesicular transport |
| Regulation of RUNX2 expression and activity | Cell signaling |

| | |
|---|---|
| Folding of actin by CCT/TriC | Cytoskeleton & Rho GTPase signaling |
| Deadenylation-dependent mRNA decay | Protein & RNA homeostasis |
| Transcription of E2F targets under negative control by DREAM complex | Cell cycle & mitosis |
| Resolution of Sister Chromatid Cohesion | Cell cycle & mitosis |
| MHC class II antigen presentation | Immune response & host–pathogen interactions |
| Degradation of AXIN | Cell signaling |
| C-type lectin receptors (CLRs) | Immune response & host–pathogen interactions |
| Negative regulation of NOTCH4 signaling | Cell signaling |
| Regulation of APC/C activators between G1/S and early anaphase | Cell cycle & mitosis |
| Translesion synthesis by POLI | DNA replication & repair |
| Homologous DNA Pairing and Strand Exchange | DNA replication & repair |
| Hedgehog 'on' state | Cell signaling |
| SARS-CoV Infections | Immune response & host–pathogen interactions |
| Defective CFTR causes cystic fibrosis | Other |
| PELO:HBS1L and ABCE1 dissociate a ribosome on a non-stop mRNA | Protein & RNA homeostasis |
| Assembly of the pre-replicative complex | DNA replication & repair |
| Recruitment of NuMA to mitotic centrosomes | Cell cycle & mitosis |
| DNA Damage Bypass | DNA replication & repair |

| | |
|---|---|
| SPOP-mediated proteasomal degradation of PD-L1(CD274) | Immune response & host–pathogen interactions |
| APC/C:Cdh1 mediated degradation of Cdc20 and other APC/C:Cdh1 targeted proteins in late mitosis/early G1 | Cell cycle & mitosis |
| Gene Silencing by RNA | Protein & RNA homeostasis |
| Processive synthesis on the C-strand of the telomere | DNA replication & repair |
| Estrogen-dependent gene expression | Cell signaling |
| Transcriptional Regulation by TP53 | Cell signaling |
| Vif-mediated degradation of APOBEC3G | Immune response & host–pathogen interactions |
| Stabilization of p53 | Cell signaling |
| Activation of the mRNA upon binding of the cap-binding complex and eIFs, and subsequent binding to 43S | Protein & RNA homeostasis |
| Eukaryotic Translation Termination | Protein & RNA homeostasis |
| GSK3B and BTRC:CUL1-mediated-degradation of NFE2L2 | Cell signaling |
| COPI-mediated anterograde transport | Membrane trafficking & vesicular transport |
| Prefoldin mediated transfer of substrate to CCT/TriC | Protein & RNA homeostasis |
| Fc epsilon receptor (FCERI) signaling | Immune response & host–pathogen interactions |
| HIV Infection | Immune response & host–pathogen interactions |
| DNA Replication Pre-Initiation | DNA replication & repair |

| | |
|---|---|
| Regulation of mRNA stability by proteins that bind AU-rich elements | Protein & RNA homeostasis |
| Formation of paraxial mesoderm | Other |
| Ribosome-associated quality control | Protein & RNA homeostasis |
| Neddylation | Ubiquitin-mediated protein degradation |
| Diseases of DNA Double-Strand Break Repair | DNA replication & repair |
| FGFR2 mutant receptor activation | Cell signaling |
| Leading Strand Synthesis | DNA replication & repair |
| Mitotic Anaphase | Cell cycle & mitosis |
| SCF(Skp2)-mediated degradation of p27/p21 | Cell cycle & mitosis |
| Processive synthesis on the lagging strand | DNA replication & repair |
| ATP-dependent chromatin remodelers | Protein & RNA homeostasis |
| Mitotic Metaphase and Anaphase | Cell cycle & mitosis |
| The role of GTSE1 in G2/M progression after G2 checkpoint | Cell cycle & mitosis |
| HDR through Homologous Recombination (HRR) or Single Strand Annealing (SSA) | DNA replication & repair |
| RNA Pol II CTD phosphorylation and interaction with CE during HIV infection | Immune response & host–pathogen interactions |
| Cytokine Signaling in Immune system | Immune response & host–pathogen interactions |
| Nervous system development | Other |

| | |
|---|---|
| ISG15 antiviral mechanism | Immune response & host–pathogen interactions |
| Eukaryotic Translation Elongation | Protein & RNA homeostasis |
| DNA Repair | DNA replication & repair |
| mRNA Polyadenylation | Protein & RNA homeostasis |
| Regulation of expression of SLITs and ROBOs | Cell signaling |
| CLEC7A (Dectin-1) signaling | Immune response & host–pathogen interactions |
| Activation of NF-kappaB in B cells | Immune response & host–pathogen interactions |
| FCERI mediated NF-kB activation | Immune response & host–pathogen interactions |
| Signaling by Interleukins | Immune response & host–pathogen interactions |
| Mitochondrial unfolded protein response (UPRmt) | Protein & RNA homeostasis |
| Export of Viral Ribonucleoproteins from Nucleus | Immune response & host–pathogen interactions |
| Regulation of RAS by GAPs | Cell signaling |
| Signaling by TGF-beta Receptor Complex | Cell signaling |
| FBXL7 down-regulates AURKA during mitotic entry and in early mitosis | Cell cycle & mitosis |
| RNA Polymerase II Promoter Escape | Protein & RNA homeostasis |
| Dual Incision in GG-NER | DNA replication & repair |
| Regulation of ornithine decarboxylase (ODC) | Cell signaling |

| | |
|---|---|
| p53-Dependent G1/S DNA damage checkpoint | DNA replication & repair |
| Regulation of T cell activation by CD28 family | Immune response & host–pathogen interactions |
| Oxygen-dependent proline hydroxylation of Hypoxia-inducible Factor Alpha | Cell signaling |
| Cilium Assembly | Membrane trafficking & vesicular transport |
| GLI3 is processed to GLI3R by the proteasome | Cell signaling |
| Antigen processing-Cross presentation | Immune response & host–pathogen interactions |
| Plasma lipoprotein assembly, remodeling, and clearance | Other |
| Macroautophagy | Autophagy |
| Translesion synthesis by POLK | DNA replication & repair |
| SARS-CoV-1-host interactions | Immune response & host–pathogen interactions |
| APC/C:Cdc20 mediated degradation of mitotic proteins | Cell cycle & mitosis |
| GTP hydrolysis and joining of the 60S ribosomal subunit | Protein & RNA homeostasis |
| Inactivation of CSF3 (G-CSF) signaling | Immune response & host–pathogen interactions |
| Downregulation of TGF-beta receptor signaling | Cell signaling |
| PTK6 Regulates RTKs and Their Effectors AKT1 and DOK1 | Cell signaling |
| Separation of Sister Chromatids | Cell cycle & mitosis |
| E3 ubiquitin ligases ubiquitinate target proteins | Ubiquitin-mediated protein degradation |

| | |
|---|---|
| Translesion Synthesis by POLH | DNA replication & repair |
| ABC transporter disorders | Other |
| CDK-mediated phosphorylation and removal of Cdc6 | Cell cycle & mitosis |
| Telomere C-strand synthesis initiation | DNA replication & repair |
| Degradation of DVL | Cell signaling |
| Activation of ATR in response to replication stress | DNA replication & repair |
| Cellular response to hypoxia | Cell signaling |
| ER to Golgi Anterograde Transport | Membrane trafficking & vesicular transport |
| Gap-filling DNA repair synthesis and ligation in GG-NER | DNA replication & repair |
| Activation of anterior HOX genes in hindbrain development during early embryogenesis | Other |
| Regulation of PTEN localization | Cell signaling |
| MTOR signalling | Cell signaling |
| Co-inhibition by PD-1 | Immune response & host–pathogen interactions |
| SRC activates STAT3 in a quantitative manner, through Cadherin-11 (CDH11), RAC1 and gp130 (IL6ST) | Cell signaling |
| Peptide chain elongation | Protein & RNA homeostasis |
| HIV Life Cycle | Immune response & host–pathogen interactions |
| NIK–>noncanonical NF-kB signaling | Immune response & host–pathogen interactions |
| mRNA Capping | Protein & RNA homeostasis |

| | |
|---|---|
| Inhibition of the proteolytic activity of APC/C required for the onset of anaphase by mitotic spindle checkpoint components | Cell cycle & mitosis |
| Synthesis of DNA | DNA replication & repair |
| Disorders of transmembrane transporters | Other |
| MAPK family signaling cascades | Cell signaling |
| Cell Cycle | Cell cycle & mitosis |
| Regulation of PLK1 Activity at G2/M Transition | Cell cycle & mitosis |
| Apoptosis | Other |
| Selective autophagy | Autophagy |
| RNA polymerase II transcribes snRNA genes | Protein & RNA homeostasis |
| Regulation of TP53 Expression and Degradation | Cell signaling |
| Formation of RNA Pol II elongation complex | Protein & RNA homeostasis |
| Gap-filling DNA repair synthesis and ligation in TC-NER | DNA replication & repair |
| Autophagy | Autophagy |
| Cross-presentation of soluble exogenous antigens (endosomes) | Immune response & host–pathogen interactions |
| Metabolism of polyamines | Other |
| Intraflagellar transport | Membrane trafficking & vesicular transport |
| TICAM1, RIP1-mediated IKK complex recruitment | Immune response & host–pathogen interactions |
| Processing of Capped Intron-Containing Pre-mRNA | Protein & RNA homeostasis |
| RNA Polymerase II Transcription Elongation | Protein & RNA homeostasis |

| | |
|---|---|
| Positive epigenetic regulation of rRNA expression | Protein & RNA homeostasis |
| Translation | Protein & RNA homeostasis |
| PTEN Regulation | Cell signaling |
| HDR through Single Strand Annealing (SSA) | DNA replication & repair |
| Activation of HOX genes during differentiation | Cell signaling |
| Dectin-1 mediated noncanonical NF-kB signaling | Immune response & host–pathogen interactions |
| Mitotic G2-G2/M phases | Cell cycle & mitosis |
| Vesicle-mediated transport | Membrane trafficking & vesicular transport |
| Cdc20:Phospho-APC/C mediated degradation of Cyclin A | Cell cycle & mitosis |
| Inactivation of APC/C via direct inhibition of the APC/C complex | Cell cycle & mitosis |
| KEAP1-NFE2L2 pathway | Cell signaling |
| Selenoamino acid metabolism | Other |
| Metabolism of amino acids and derivatives | Other |
| Regulation of PD-L1(CD274) Post-translational modification | Immune response & host–pathogen interactions |
| Innate Immune System | Immune response & host–pathogen interactions |
| Formation of the Early Elongation Complex | Protein & RNA homeostasis |
| Regulation of RUNX3 expression and activity | Cell signaling |
| FLT3 signaling by CBL mutants | Cell signaling |
| Diseases of programmed cell death | Other |
| Translesion synthesis by REV1 | DNA replication & repair |

| | |
|---|---|
| APC:Cdc20 mediated degradation of cell cycle proteins prior to satisfation of the cell cycle checkpoint | Cell cycle & mitosis |
| Antigen processing: Ubiquitination & Proteasome degradation | Immune response & host–pathogen interactions |
| Chromosome Maintenance | DNA replication & repair |
| Signaling by Rho GTPases | Cytoskeleton & Rho GTPase signaling |
| Resolution of D-loop Structures through Holliday Junction Intermediates | DNA replication & repair |
| Influenza Viral RNA Transcription and Replication | Immune response & host–pathogen interactions |
| InlA-mediated entry of Listeria monocytogenes into host cells | Immune response & host–pathogen interactions |
| Mismatch Repair | DNA replication & repair |
| Circadian clock | Other |
| Impaired BRCA2 binding to PALB2 | DNA replication & repair |
| Tat-mediated HIV elongation arrest and recovery | Immune response & host–pathogen interactions |
| IRAK1 recruits IKK complex | Immune response & host–pathogen interactions |
| Neutrophil degranulation | Immune response & host–pathogen interactions |
| Ubiquitin-dependent degradation of Cyclin D | Cell cycle & mitosis |
| PINK1-PRKN Mediated Mitophagy | Autophagy |

| | |
|---|---|
| Telomere Maintenance | DNA replication & repair |
| Amino acids regulate mTORC1 | Cell signaling |
| FGFR2 alternative splicing | Protein & RNA homeostasis |
| DNA Double-Strand Break Repair | DNA replication & repair |
| TCR signaling | Immune response & host–pathogen interactions |
| IKK complex recruitment mediated by RIP1 | Immune response & host–pathogen interactions |
| Removal of the Flap Intermediate | DNA replication & repair |
| Metabolism of RNA | Protein & RNA homeostasis |
| Regulation of PD-L1(CD274) expression | Immune response & host–pathogen interactions |
| Myoclonic epilepsy of Lafora | Other |
| Dengue Virus-Host Interactions | Immune response & host–pathogen interactions |
| Adherens junctions interactions | Cytoskeleton & Rho GTPase signaling |
| SARS-CoV-2-host interactions | Immune response & host–pathogen interactions |
| Recognition of DNA damage by PCNA-containing replication complex | DNA replication & repair |
| Activation of the pre-replicative complex | DNA replication & repair |

| | |
|---|---|
| RHO GTPases Activate Formins | Cytoskeleton & Rho GTPase signaling |
| Adaptive Immune System | Immune response & host–pathogen interactions |
| G2/M Transition | Cell cycle & mitosis |
| TCF dependent signaling in response to WNT | Cell signaling |
| Maturation of protein E | Immune response & host–pathogen interactions |
| Class I MHC mediated antigen processing & presentation | Immune response & host–pathogen interactions |
| EML4 and NUDC in mitotic spindle formation | Cell cycle & mitosis |
| tRNA Aminoacylation | Protein & RNA homeostasis |
| RHO GTPases Activate WASPs and WAVEs | Cytoskeleton & Rho GTPase signaling |
| p53-Independent G1/S DNA Damage Checkpoint | DNA replication & repair |
| Selenocysteine synthesis | Other |
| activated TAK1 mediates p38 MAPK activation | Cell signaling |
| GSK3B-mediated proteasomal degradation of PD-L1(CD274) | Immune response & host–pathogen interactions |
| Nucleotide Excision Repair | DNA replication & repair |
| Transport of the SLBP independent Mature mRNA | Protein & RNA homeostasis |
| Plasma lipoprotein clearance | Other |

| | |
|---|---|
| COPI-dependent Golgi-to-ER retrograde traffic | Membrane trafficking & vesicular transport |
| MicroRNA (miRNA) biogenesis | Protein & RNA homeostasis |
| Transport of small molecules | Membrane trafficking & vesicular transport |
| rRNA processing | Protein & RNA homeostasis |
| SARS-CoV-2 activates/modulates innate and adaptive immune responses | Immune response & host–pathogen interactions |
| Hh mutants abrogate ligand secretion | Cell signaling |
| PCNA-Dependent Long Patch Base Excision Repair | DNA replication & repair |
| TRAF6 mediated IRF7 activation in TLR7/8 or 9 signaling | Immune response & host–pathogen interactions |
| ABC-family protein mediated transport | Membrane trafficking & vesicular transport |
| Somitogenesis | Other |
| Gastrulation | Other |
| Homology Directed Repair | DNA replication & repair |
| PIWI-interacting RNA (piRNA) biogenesis | Protein & RNA homeostasis |
| Dual incision in TC-NER | DNA replication & repair |
| SRP-dependent cotranslational protein targeting to membrane | Protein & RNA homeostasis |
| DNA strand elongation | DNA replication & repair |
| Regulation of TP53 Activity through Phosphorylation | Cell signaling |
| Degradation of beta-catenin by the destruction complex | Cell signaling |

| | |
|---|---|
| S Phase | Cell cycle & mitosis |
| Translation initiation complex formation | Protein & RNA homeostasis |
| G1/S DNA Damage Checkpoints | DNA replication & repair |
| G2/M DNA damage checkpoint | DNA replication & repair |
| Presynaptic phase of homologous DNA pairing and strand exchange | DNA replication & repair |
| Interleukin-1 signaling | Immune response & host–pathogen interactions |
| Cellular responses to stress | Other |
| Modulation by Mtb of host immune system | Immune response & host–pathogen interactions |
| RHO GTPase Effectors | Cytoskeleton & Rho GTPase signaling |
| Diseases of DNA repair | DNA replication & repair |
| Polymerase switching | DNA replication & repair |
| Ub-specific processing proteases | Ubiquitin-mediated protein degradation |
| Signaling by Rho GTPases, Miro GTPases and RHOBTB3 | Cytoskeleton & Rho GTPase signaling |
| MAPK1/MAPK3 signaling | Cell signaling |
| Assembly of the 9+0 primary cilium | Membrane trafficking & vesicular transport |

| | |
|---|---|
| Transcription of the HIV genome | Immune response & host–pathogen interactions |
| ER-Phagosome pathway | Membrane trafficking & vesicular transport |
| M Phase | Cell cycle & mitosis |
| Downregulation of ERBB4 signaling | Cell signaling |
| Degradation of GLI1 by the proteasome | Cell signaling |
| Formation of TC-NER Pre-Incision Complex | DNA replication & repair |
| Polymerase switching on the C-strand of the telomere | DNA replication & repair |
| Cellular responses to stimuli | Other |
| RNA Polymerase II Transcription Pre-Initiation And Promoter Opening | Protein & RNA homeostasis |
| Switching of origins to a post-replicative state | DNA replication & repair |
| SARS-CoV-1 Infection | Immune response & host–pathogen interactions |
| Antigen processing: Ub, ATP-independent proteasomal degradation | Immune response & host–pathogen interactions |
| Antimicrobial mechanism of IFN-stimulated genes | Immune response & host–pathogen interactions |
| Protein folding | Protein & RNA homeostasis |
| Resolution of D-loop Structures through Synthesis-Dependent Strand Annealing (SDSA) | DNA replication & repair |
| Protein ubiquitination | Ubiquitin-mediated protein degradation |

| | |
|---|---|
| Interferon Signaling | Immune response & host–pathogen interactions |
| Cap-dependent Translation Initiation | Protein & RNA homeostasis |
| Mitotic G1 phase and G1/S transition | Cell cycle & mitosis |
| Dengue Virus Genome Translation and Replication | Immune response & host–pathogen interactions |
| Transcriptional regulation by RUNX1 | Cell signaling |
| Late Phase of HIV Life Cycle | Immune response & host–pathogen interactions |
| Transport of Mature Transcript to Cytoplasm | Protein & RNA homeostasis |
| Organelle biogenesis and maintenance | Other |
| Defective homologous recombination repair (HRR) due to BRCA2 loss of function | DNA replication & repair |
| Prevention of phagosomal-lysosomal fusion | Immune response & host–pathogen interactions |
| Transport to the Golgi and subsequent modification | Membrane trafficking & vesicular transport |
| Signaling by EGFR in Cancer | Cell signaling |
| Activation of APC/C and APC/C:Cdc20 mediated degradation of mitotic proteins | Cell cycle & mitosis |
| Transport of the SLBP Dependant Mature mRNA | Protein & RNA homeostasis |
| Regulation of Apoptosis | Cell signaling |
| SARS-CoV-2 modulates host translation machinery | Immune response & host–pathogen interactions |

| | |
|---|---|
| Influenza Infection | Immune response & host–pathogen interactions |
| Cell Cycle Checkpoints | Cell cycle & mitosis |
| Resolution of D-Loop Structures | DNA replication & repair |
| Cell junction organization | Cytoskeleton & Rho GTPase signaling |
| Disease | Other |
| Interleukin-1 family signaling | Immune response & host–pathogen interactions |
| Metabolism of non-coding RNA | Protein & RNA homeostasis |
| Cyclin E associated events during G1/S transition | Cell cycle & mitosis |
| Formation of tubulin folding intermediates by CCT/TriC | Cytoskeleton & Rho GTPase signaling |
| MITF-M-regulated melanocyte development | Cell signaling |
| AMPK-induced ERAD and lysosome mediated degradation of PD-L1(CD274) | Immune response & host–pathogen interactions |
| Viral Infection Pathways | Immune response & host–pathogen interactions |
| Association of TriC/CCT with target proteins during biosynthesis | Protein & RNA homeostasis |
| Orc1 removal from chromatin | DNA replication & repair |
| Cargo recognition for clathrin-mediated endocytosis | Membrane trafficking & vesicular transport |

| | |
|---|---|
| UCH proteinases | Ubiquitin-mediated protein degradation |

### Effect of refinement on functional landscape of top-ranked genes

**Table S10: Comparison of overrepresentation analysis (ORA) results for different intensity regimes.**

| Category | average_minus_log10(Benjamini) | n_terms | genes_fraction | ORA/intensity regime |
|---|---|---|---|---|
| Cell cycle & mitosis | 4.629 | 36 | 0.084 | early_low_unrefined |
| DNA replication & repair | 4.463 | 12 | 0.06 | early_low_unrefined |
| Cytoskeleton & Rho GTPase signaling | 5.840 | 6 | 0.224 | early_low_unrefined |
| Membrane trafficking & vesicular transport | 2.415 | 6 | 0.076 | early_low_unrefined |
| Immune response & host–pathogen interactions | 7.966 | 51 | 0.376 | early_low_unrefined |
| Protein & RNA homeostasis | 23.244 | 45 | 0.472 | early_low_unrefined |
| Ubiquitin-mediated protein degradation | 4.501 | 9 | 0.06 | early_low_unrefined |
| Autophagy | 36.020 | 1 | 0.164 | early_low_unrefined |
| Cell signaling | 5.783 | 59 | 0.264 | early_low_unrefined |
| Other | 12.664 | 21 | 0.528 | early_low_unrefined |
| Cell cycle & mitosis | 17.923 | 52 | 0.284 | early_low_refined |
| DNA replication & repair | 7.571 | 55 | 0.26 | early_low_refined |

| | | | | |
|---|---|---|---|---|
| Cytoskeleton & Rho GTPase signaling | 7.228 | 11 | 0.252 | early_low_ refined |
| Membrane trafficking & vesicular transport | 5.168 | 17 | 0.272 | early_low_ refined |
| Immune response & host–pathogen interactions | 9.622 | 74 | 0.364 | early_low_ refined |
| Protein & RNA homeostasis | 5.835 | 45 | 0.544 | early_low_ refined |
| Ubiquitin-mediated protein degradation | 15.078 | 14 | 0.192 | early_low_ refined |
| Autophagy | 2.258 | 8 | 0.072 | early_low_ refined |
| Cell signaling | 12.509 | 74 | 0.372 | early_low_ refined |
| Other | 11.437 | 23 | 0.408 | early_low_ refined |
| Cell cycle & mitosis | 1.750 | 2 | 0.1 | early_high_ unrefined |
| DNA replication & repair | 1.797 | 3 | 0.036 | early_high_ unrefined |
| Cytoskeleton & Rho GTPase signaling | 0.000 | 0 | 0.0 | early_high_ unrefined |
| Membrane trafficking & vesicular transport | 0.000 | 0 | 0.0 | early_high_ unrefined |
| Immune response & host–pathogen interactions | 1.943 | 19 | 0.148 | early_high_ unrefined |
| Protein & RNA homeostasis | 1.810 | 18 | 0.052 | early_high_ unrefined |

| | | | | |
|---|---|---|---|---|
| Ubiquitin-mediated protein degradation | 0.000 | 0 | 0.0 | early_high_unrefined |
| Autophagy | 0.000 | 0 | 0.0 | early_high_unrefined |
| Cell signaling | 1.631 | 4 | 0.036 | early_high_unrefined |
| Other | 1.467 | 3 | 0.2 | early_high_unrefined |
| Cell cycle & mitosis | 6.146 | 26 | 0.24 | early_high_refined |
| DNA replication & repair | 4.460 | 57 | 0.192 | early_high_refined |
| Cytoskeleton & Rho GTPase signaling | 3.884 | 7 | 0.132 | early_high_refined |
| Membrane trafficking & vesicular transport | 2.858 | 10 | 0.148 | early_high_refined |
| Immune response & host–pathogen interactions | 2.359 | 8 | 0.292 | early_high_refined |
| Protein & RNA homeostasis | 3.649 | 7 | 0.26 | early_high_refined |
| Ubiquitin-mediated protein degradation | 1.461 | 1 | 0.048 | early_high_refined |
| Autophagy | 2.172 | 4 | 0.044 | early_high_refined |
| Cell signaling | 3.234 | 4 | 0.12 | early_high_refined |
| Other | 2.777 | 6 | 0.356 | early_high_refined |
| Cell cycle & mitosis | 8.912 | 35 | 0.092 | late_low_unrefined |

| | | | | |
|---|---|---|---|---|
| DNA replication & repair | 8.913 | 10 | 0.06 | late_low_ unrefined |
| Cytoskeleton & Rho GTPase signaling | 6.964 | 6 | 0.192 | late_low_ unrefined |
| Membrane trafficking & vesicular transport | 7.604 | 3 | 0.1 | late_low_ unrefined |
| Immune response & host–pathogen interactions | 9.611 | 44 | 0.3 | late_low_ unrefined |
| Protein & RNA homeostasis | 22.215 | 35 | 0.396 | late_low_ unrefined |
| Ubiquitin-mediated protein degradation | 9.574 | 9 | 0.088 | late_low_ unrefined |
| Autophagy | 26.330 | 1 | 0.132 | late_low_ unrefined |
| Cell signaling | 10.432 | 47 | 0.22 | late_low_ unrefined |
| Other | 12.343 | 21 | 0.408 | late_low_ unrefined |
| Cell cycle & mitosis | 23.620 | 47 | 0.352 | late_low_ refined |
| DNA replication & repair | 9.829 | 77 | 0.328 | late_low_ refined |
| Cytoskeleton & Rho GTPase signaling | 7.246 | 14 | 0.244 | late_low_ refined |
| Membrane trafficking & vesicular transport | 5.436 | 13 | 0.264 | late_low_ refined |
| Immune response & host–pathogen interactions | 15.330 | 50 | 0.348 | late_low_ refined |

| | | | | |
|---|---|---|---|---|
| Protein & RNA homeostasis | 9.015 | 25 | 0.54 | late_low_ refined |
| Ubiquitin-mediated protein degradation | 20.484 | 12 | 0.22 | late_low_ refined |
| Autophagy | 2.511 | 6 | 0.048 | late_low_ refined |
| Cell signaling | 19.360 | 54 | 0.372 | late_low_ refined |
| Other | 14.470 | 20 | 0.4 | late_low_ refined |
| Cell cycle & mitosis | 0.000 | 0 | 0.0 | late_high_ unrefined |
| DNA replication & repair | 0.000 | 0 | 0.0 | late_high_ unrefined |
| Cytoskeleton & Rho GTPase signaling | 0.000 | 0 | 0.0 | late_high_ unrefined |
| Membrane trafficking & vesicular transport | 0.000 | 0 | 0.0 | late_high_ unrefined |
| Immune response & host–pathogen interactions | 0.000 | 0 | 0.0 | late_high_ unrefined |
| Protein & RNA homeostasis | 0.000 | 0 | 0.0 | late_high_ unrefined |
| Ubiquitin-mediated protein degradation | 0.000 | 0 | 0.0 | late_high_ unrefined |
| Autophagy | 0.000 | 0 | 0.0 | late_high_ unrefined |
| Cell signaling | 0.000 | 0 | 0.0 | late_high_ unrefined |
| Other | 0.000 | 0 | 0.0 | late_high_ unrefined |

| | | | | |
|---|---|---|---|---|
| Cell cycle & mitosis | 1.605 | 4 | 0.104 | late_high_refined |
| DNA replication & repair | 2.137 | 1 | 0.072 | late_high_refined |
| Cytoskeleton & Rho GTPase signaling | 1.329 | 2 | 0.1 | late_high_refined |
| Membrane trafficking & vesicular transport | 0.000 | 0 | 0.0 | late_high_refined |
| Immune response & host–pathogen interactions | 1.941 | 2 | 0.24 | late_high_refined |
| Protein & RNA homeostasis | 0.000 | 0 | 0.0 | late_high_refined |
| Ubiquitin-mediated protein degradation | 0.000 | 0 | 0.0 | late_high_refined |
| Autophagy | 2.036 | 2 | 0.048 | late_high_refined |
| Cell signaling | 0.000 | 0 | 0.0 | late_high_refined |
| Other | 0.000 | 0 | 0.0 | late_high_refined |

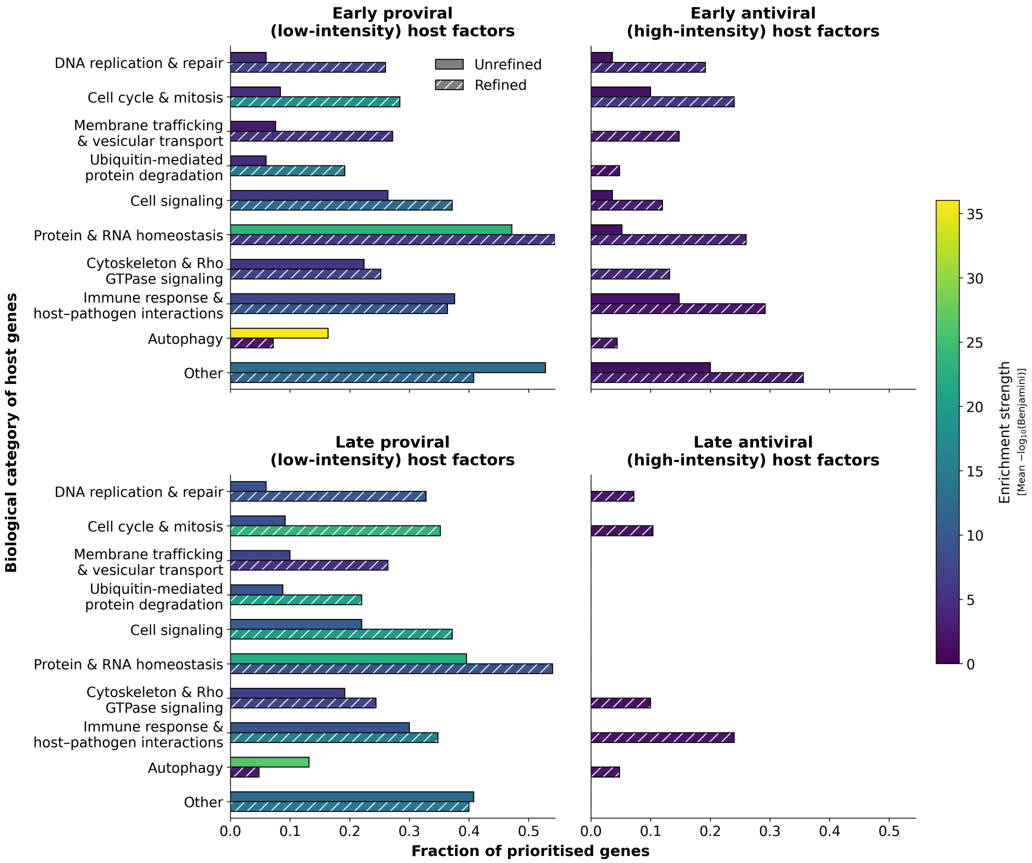


**Figure S10: Effect of PPI-informed confidence weighting on the functional landscape of top-ranked genes.** Horizontal bar length indicates the fraction of top-ranked genes assigned to each biological category, whereas bar colour indicates enrichment strength, quantified as the mean -log10(Benjamini) value of Reactome pathway terms assigned to the respective category.

### References


[1] Diego E Alvarez and Hervé Agaisse. Casein kinase 2 regulates vaccinia virus actin tail formation. *Virology*, 423(2):143–151, 2012.

[2] Diego E Alvarez and Hervé Agaisse. The formin fhod1 and the small gtpase rac1 promote vaccinia virus actin–based motility. *Journal of Cell Biology*, 202(7):1075–1090, 2013.

[3] Anderson A Andrade, Patricia NG Silva, Anna CTC Pereira, Lirlândia P De Sousa, Paulo CP Ferreira, Ricardo T Gazzinelli, Erna G Kroon, Catherine Ropert, and Cláudio A Bonjardim. The vaccinia virus-stimulated mitogen-activated protein kinase (mapk) pathway is required for virus multiplication. *Biochemical Journal*, 381(2):437–446, 2004.

[4] Angika Basant and Michael Way. The relative binding position of nck and grb2 adaptors impacts actin-based motility of vaccinia virus. *Elife*, 11:e74655, 2022.

[5] Philippa M Beard, Samantha J Griffiths, Orland Gonzalez, Ismar R Haga, Tali Pechenick Jowers, Danielle K Reynolds, Jan Wildenhain, Hille Tekotte, Manfred Auer, Mike Tyers, et al. A loss of function analysis of host factors influencing vaccinia virus replication by rna interference. *PLoS One*, 9(6):e98431, 2014.

[6] Martina Bécares, Manuel Albert, Céline Tárrega, Rocío Coloma, Michela Falqui, Emma K Luhmann, Lilliana Radoshevich, and Susana Guerra. Isg15 is required for the dissemination of vaccinia virus extracellular virions. *Microbiology spectrum*, 11(3):e04508–22, 2023.

[7] Andrew Bowie, Endre Kiss-Toth, Julian A Symons, Geoffrey L Smith, Steven K Dower, and Luke AJ O'Neill. A46r and a52r from vaccinia virus are antagonists of host il-1 and toll-like receptor signaling. *Proceedings of the National Academy of Sciences*, 97(18):10162–10167, 2000.

[8] YoungSik Cho, Sreerupa Challa, David Moquin, Ryan Genga, Tathagat Dutta Ray, Melissa Guildford, and Francis Ka-Ming Chan. Phosphorylation-driven assembly of the rip1-rip3 complex regulates programmed necrosis and virus-induced inflammation. *Cell*, 137(6):1112–1123, 2009.

[9] Sally Cudmore, Pascale Cossart, Gareth Griffiths, and Michael Way. Actin-based motility of vaccinia virus. *Nature*, 378(6557):636–638, 1995.

[10] Rachel David. Propelling vaccinia virus to the neighbours. *Nature Reviews Microbiology*, 11(9):595–595, 2013.

[11] José C de Magalhaes, Anderson A Andrade, Patrícia NG Silva, Lirlândia P Sousa, Catherine Ropert, Paulo CP Ferreira, Erna G Kroon, Ricardo T Gazzinelli, and Cláudio A Bonjardim. A mitogenic signal triggered at an early stage of vaccinia virus infection: implication of mek/erk and protein kinase a in virus multiplication. *Journal of Biological Chemistry*, 276(42):38353–38360, 2001.

[12] Pragyesh Dhungel, Fernando M Cantu, Joshua A Molina, and Zhilong Yang. Vaccinia virus as a master of host shutoff induction: targeting processes of the central dogma and beyond. *Pathogens*, 9(5):400, 2020.

[13] Deborah A Eppstein, Y Vivienne Marsh, Alain B Schreiber, Sherry R Newman, George J Todaro, and John J Nestor Jr. Epidermal growth factor receptor occupancy inhibits vaccinia virus infection. *Nature*, 318(6047):663–665, 1985.

[14] Iliana Georgana, Simon R Scutts, Chen Gao, Yongxu Lu, Alice A Torres, Hongwei Ren, Edward Emmott, Jinghao Men, Keefe Oei, and Geoffrey L Smith. Filamin b restricts vaccinia virus spread and is targeted by vaccinia virus protein c4. *Journal of virology*, 98(3):e01485–23, 2024.

[15] Motti Gerlic, Benjamin Faustin, Antonio Postigo, Eric Chi-Wang Yu, Martina Proell, Naran Gombosuren, Maryla Krajewska, Rachel Flynn, Michael Croft, Michael Way, et al. Vaccinia virus f1l protein promotes virulence by inhibiting inflammasome activation. *Proceedings of the National Academy of Sciences*, 110(19):7808–7813, 2013.

[16] James Harris and Natalie A Borg. The multifaceted roles of nlrp3-modulating proteins in virus infection. *Frontiers in Immunology*, 13:987453, 2022.

[17] Crispin T. Hiley, Louisa S. Chard, Rathi Gangeswaran, James R. Tysome, Arnaud Briat, Nicholas R. Lemoine, and Yaohe Wang. Vascular endothelial growth factor a promotes vaccinia virus entry into host

cells via activation of the akt pathway. *Journal of virology*, 87(5):2781, 2013.

[18] Cheng-Yen Huang, Tsai-Yi Lu, Chi-Horng Bair, Yuan-Shau Chang, Jeng-Kuan Jwo, and Wen Chang. A novel cellular protein, vpef, facilitates vaccinia virus penetration into hela cells through fluid phase endocytosis. *Journal of virology*, 82(16):7988–7999, 2008.

[19] Ashley C Humphries, Sara K Donnelly, and Michael Way. Cdc42 and the rho gef intersectin-1 collaborate with nck to promote n-wasp-dependent actin polymerisation. *Journal of cell science*, 127(3):673–685, 2014.

[20] Jan-Jong Hung, Che-Sheng Chung, and Wen Chang. Molecular chaperone hsp90 is important for vaccinia virus growth in cells. *Journal of virology*, 76(3):1379–1390, 2002.

[21] Moona Huttunen, Jerzy Samolej, Robert J Evans, Artur Yakimovich, Ian J White, Janos Kriston-Vizi, Juan Martin-Serrano, Wesley I Sundquist, Eva-Maria Frickel, and Jason Mercer. Vaccinia virus hijacks escrt-mediated multivesicular body formation for virus egress. *Life Science Alliance*, 4(8):e202000910, 2021.

[22] Nouhou Ibrahim, April Wicklund, and Matthew S Wiebe. Molecular characterization of the host defense activity of the barrier to autointegration factor against vaccinia virus. *Journal of virology*, 85(22):11588–11600, 2011.

[23] Roza Izmailyan, Jye-Chian Hsao, Che-Sheng Chung, Chein-Hung Chen, Paul Wei-Che Hsu, Chung-Lin Liao, and Wen Chang. Integrin $\beta 1$ mediates vaccinia virus entry through activation of pi3k/akt signaling. *Journal of virology*, 86(12):6677–6687, 2012.

[24] Satish Jindal and RICHARD A Young. Vaccinia virus infection induces a stress response that leads to association of hsp70 with viral proteins. *Journal of Virology*, 66(9):5357–5362, 1992.

[25] Tali Pechenick Jowers, Rebecca J Featherstone, Danielle K Reynolds, Helen K Brown, John James, Alan Prescott, Ismar R Haga, and Philippa M Beard. Rab1a promotes vaccinia virus replication by facilitating the production of intracellular enveloped virions. *Virology*, 475:66–73, 2015.

[26] George C Katsafanas and Bernard Moss. Vaccinia virus intermediate stage transcription is complemented by ras-gtpase-activating protein sh3 domain-binding protein (g3bp) and cytoplasmic activation/proliferation-associated protein (p137) individually or as a heterodimer. *Journal of Biological Chemistry*, 279(50):52210–52217, 2004.

[27] Alexander Kuehnl, Agnes Musiol, Carsten A Raabe, and Ursula Rescher. Emerging functions as host cell factors–an encyclopedia of annexin-pathogen interactions. *Biological Chemistry*, 397(10):949–959, 2016.

[28] Flavia Leite and Michael Way. The role of signalling and the cytoskeleton during vaccinia virus egress. *Virus research*, 209:87–99, 2015.

[29] Chang Li, Shouwen Du, Mingyao Tian, Yuhang Wang, Jieying Bai, Peng Tan, Wei Liu, Ronglan Yin, Maopeng Wang, Ying Jiang, et al. The host restriction factor interferon-inducible transmembrane protein 3 inhibits vaccinia virus infection. *Frontiers in immunology*, 9:228, 2018.

[30] Ye Li, Leiliang Zhang, and Youyang Ke. Cellular interactome analysis of vaccinia virus k7 protein identifies three transport machineries as binding partners for k7. *Virus Genes*, 53(6):814–822, 2017.

[31] Yu Li, Staffan Grenklo, Theresa Higgins, and Roger Karlsson. The profilin: actin complex localizes to sites of dynamic actin polymerization at the leading edge of migrating cells and pathogen-induced actin tails. *European journal of cell biology*, 87(11):893–904, 2008.

[32] Alexandria C Linville, Amber B Rico, Helena Teague, Lucy E Binsted, Geoffrey L Smith, Jonas D Albarnaz, and Matthew S Wiebe. Dysregulation of cellular vrk1, baf, and innate immune signaling by the vaccinia virus b12 pseudokinase. *Journal of Virology*, 96(11):e00398–22, 2022.

[33] Jia Liu and Grant McFadden. Samd9 is an innate antiviral host factor with stress response properties that can be antagonized by poxviruses. *Journal of virology*, 89(3):1925–1931, 2015.

[34] Ruikang Liu, Lisa R Olano, Yeva Mirzakhanyan, Paul D Gershon, and Bernard Moss. Vaccinia virus ankyrin-repeat/f-box protein targets interferon-induced ifits for proteasomal degradation. *Cell reports*, 29(4):816–828, 2019.

[35] Rutger D Luteijn, Ferdy van Diemen, Vincent A Blomen, Ingrid GJ Boer, Saravanan Manikam Sadasivam, Toin H van Kuppevelt, Ingo Drexler, Thijn R Brummelkamp, Robert Jan Lebbink, and Emmanuel J Wiertz. A genome-wide haploid genetic screen identifies heparan sulfate-associated genes and the macropinocytosis modulator tmed10 as factors supporting vaccinia virus infection. *Journal of virology*, 93(13):10–1128, 2019.

[36] Brandon A Mann, Julia He Huang, Ping Li, Hua-Chen Chang, Roger B Slee, Audrey O'Sullivan, Mathur Anita, Norman Yeh, Michael J Klemsz, Randy R Brutkiewicz, et al. Vaccinia virus blocks stat1-dependent and stat1-independent gene expression induced by type i and type ii interferons. *Journal of Interferon & Cytokine Research*, 28(6):367–380, 2008.

[37] Alejandro Matía, Maria M Lorenzo, Yolimar C Romero-Estremera, Juana M Sánchez-Puig, Angel Zaballos, and Rafael Blasco. Identification of $\beta$2 microglobulin, the product of b2m gene, as a host factor for vaccinia virus infection by genome-wide crispr genetic screens. *PLoS Pathogens*, 18(12):e1010800, 2022.

[38] Shannon McNulty, William Bornmann, Jill Schriewer, Chas Werner, Scott K Smith, Victoria A Olson, Inger K Damon, R Mark Buller, John Heuser, and Daniel Kalman. Multiple phosphatidylinositol 3-kinases regulate vaccinia virus morphogenesis. *PLoS One*, 5(5):e10884, 2010.

[39] Nathan Meade, Melvin King, Joshua Munger, and Derek Walsh. mtor dysregulation by vaccinia virus f17 controls multiple processes with varying roles in infection. *Journal of virology*, 93(15):10–1128, 2019.

[40] Jason Mercer and Ari Helenius. Vaccinia virus uses macropinocytosis and apoptotic mimicry to enter host cells. *Science*, 320(5875):531–535, 2008.

[41] Jason Mercer, Berend Snijder, Raphael Sacher, Christine Burkard, Christopher Karl Ernst Bleck, Henning Stahlberg, Lucas Pelkmans, and Ari Helenius. Rnai screening reveals proteasome-and cullin3-dependent stages in vaccinia virus infection. *Cell reports*, 2(4):1036–1047, 2012.

[42] Kouki Morizono, Yiming Xie, Tove Olafsen, Benhur Lee, Asim Dasgupta, Anna M Wu, and Irvin SY Chen. The soluble serum protein gas6

bridges virion envelope phosphatidylserine to the tam receptor tyrosine kinase axl to mediate viral entry. *Cell host & microbe*, 9(4):286–298, 2011.

[43] Theresa S Moser, Russell G Jones, Craig B Thompson, Carolyn B Coyne, and Sara Cherry. A kinome rnai screen identified ampk as promoting poxvirus entry through the control of actin dynamics. *PLoS pathogens*, 6(6):e1000954, 2010.

[44] Bianca T Hovey Nerenberg, John Taylor, Eric Bartee, Kristine Gouveia, Michele Barry, and Klaus Früh. The poxviral ring protein p28 is a ubiquitin ligase that targets ubiquitin to viral replication factories. *Journal of virology*, 79(1):597–601, 2005.

[45] Annabel T Olson, Stephanie J Child, and Adam P Geballe. Antagonism of protein kinase r by large dna viruses. *Pathogens*, 11(7):790, 2022.

[46] Mitchell A Pallett, Hongwei Ren, Rui-Yao Zhang, Simon R Scutts, Laura Gonzalez, Zihan Zhu, Carlos Maluquer de Motes, and Geoffrey L Smith. Vaccinia virus bbk e3 ligase adaptor a55 targets importin-dependent nf-$\kappa$b activation and inhibits cd8+ t-cell memory. *Journal of virology*, 93(10):10–1128, 2019.

[47] Antonio Postigo, Amy E Ramsden, Michael Howell, and Michael Way. Cytoplasmic atr activation promotes vaccinia virus genome replication. *Cell reports*, 19(5):1022–1032, 2017.

[48] Ramtin Rahbar, Thomas T Murooka, Anna A Hinek, Carole L Galligan, Antonella Sassano, Celeste Yu, Kishore Srivastava, Leonidas C Platanias, and Eleanor N Fish. Vaccinia virus activation of ccr5 invokes tyrosine phosphorylation signaling events that support virus replication. *Journal of virology*, 80(14):7245–7259, 2006.

[49] Patrick C Reading and Geoffrey L Smith. Vaccinia virus interleukin-18-binding protein promotes virulence by reducing gamma interferon production and natural killer and t-cell activity. *Journal of virology*, 77(18):9960–9968, 2003.

[50] Susan Realegeno, Lalita Priyamvada, Amrita Kumar, Jessica B Blackburn, Claire Hartloge, Andreas S Puschnik, Suryaprakash Sambhara,

Victoria A Olson, Jan E Carette, Vladimir Lupashin, et al. Conserved oligomeric golgi (cog) complex proteins facilitate orthopoxvirus entry, fusion and spread. *Viruses*, 12(7):707, 2020.

[51] Zaira Rizopoulos, Giuseppe Balistreri, Samuel Kilcher, Caroline K Martin, Mohammedyaseen Syedbasha, Ari Helenius, and Jason Mercer. Vaccinia virus infection requires maturation of macropinosomes. *Traffic*, 16(8):814–831, 2015.

[52] Susanne Roth, Andrea Rottach, Amelie S Lotz-Havla, Verena Laux, Andreas Muschaweckh, Søren W Gersting, Ania C Muntau, Karl-Peter Hopfner, Lei Jin, Katelynd Vanness, et al. Rad50-card9 interactions link cytosolic dna sensing to il-1$\beta$ production. *Nature immunology*, 15(6):538–545, 2014.

[53] Niki Scaplehorn, Anna Holmström, Violaine Moreau, Freddy Frischknecht, Inge Reckmann, and Michael Way. Grb2 and nck act cooperatively to promote actin-based motility of vaccinia virus. *Current Biology*, 12(9):740–745, 2002.

[54] Florian I Schmidt and Jason Mercer. Vaccinia virus egress: actin out with clathrin. *Cell host & microbe*, 12(3):263–265, 2012.

[55] Simon R Scutts, Stuart W Ember, Hongwei Ren, Chao Ye, Christopher A Lovejoy, Michela Mazzon, David L Veyer, Rebecca P Sumner, and Geoffrey L Smith. Dna-pk is targeted by multiple vaccinia virus proteins to inhibit dna sensing. *Cell reports*, 25(7):1953–1965, 2018.

[56] Gilad Sivan, Pinar Ormanoglu, Eugen C Buehler, Scott E Martin, and Bernard Moss. Identification of restriction factors by human genome-wide rna interference screening of viral host range mutants exemplified by discovery of samd9 and wdr6 as inhibitors of the vaccinia virus k1l-c7l- mutant. *MBio*, 6(4):10–1128, 2015.

[57] Jamária AP Soares, Flávia GG Leite, Luciana G Andrade, Alice A Torres, Lirlândia P De Sousa, Lucíola S Barcelos, Mauro M Teixeira, Paulo CP Ferreira, Erna G Kroon, Thaís Souto-Padrón, et al. Activation of the pi3k/akt pathway early during vaccinia and cowpox virus infections is required for both host survival and viral replication. *Journal of virology*, 83(13):6883–6899, 2009.

[58] Julianne Stack, Ismar R Haga, Martina Schröder, Nathan W Bartlett, Geraldine Maloney, Patrick C Reading, Katherine A Fitzgerald, Geoffrey L Smith, and Andrew G Bowie. Vaccinia virus protein a46r targets multiple toll-like–interleukin-1 receptor adaptors and contributes to virulence. *The Journal of experimental medicine*, 201(6):1007–1018, 2005.

[59] Jennifer H Stuart, Rebecca P Sumner, Yongxu Lu, Joseph S Snowden, and Geoffrey L Smith. Vaccinia virus protein c6 inhibits type i ifn signalling in the nucleus and binds to the transactivation domain of stat2. *PLoS pathogens*, 12(12):e1005955, 2016.

[60] Alan C Townsley, Andrea S Weisberg, Timothy R Wagenaar, and Bernard Moss. Vaccinia virus entry into cells via a low-ph-dependent endosomal pathway. *Journal of virology*, 80(18):8899–8908, 2006.

[61] Zoe Waibler, Martina Anzaghe, Theresa Frenz, Astrid Schwantes, Christopher Pöhlmann, Holger Ludwig, Marcos Palomo-Otero, Antonio Alcamí, Gerd Sutter, and Ulrich Kalinke. Vaccinia virus-mediated inhibition of type i interferon responses is a multifactorial process involving the soluble type i interferon receptor b18 and intracellular components. *Journal of virology*, 83(4):1563–1571, 2009.

[62] Shuqin Xu, Xin Zhang, Chentao Li, Zixuan Zhang, Xiaoqin Wang, Chenwen Wan, Peihua Wang, Yi Lv, Yingli He, Francis Ka-Ming Chan, et al. Targeting endolysosomal acidification inhibits poxvirus entry and replication. *Cell Communication and Signaling*, 24(1):129, 2026.

[63] Izabela Zaborowska, Kerstin Kellner, Michael Henry, Paula Meleady, and Derek Walsh. Recruitment of host translation initiation factor eif4g by the vaccinia virus ssdna-binding protein i3. *Virology*, 425(1):11–22, 2012.

[64] Leiliang Zhang, Stella Y Lee, Galina V Beznoussenko, Peter J Peters, Jia-Shu Yang, Hui-ya Gilbert, Abraham L Brass, Stephen J Elledge, Stuart N Isaacs, Bernard Moss, et al. A role for the host coatomer and kdel receptor in early vaccinia biogenesis. *Proceedings of the National Academy of Sciences*, 106(1):163–168, 2009.

[65] Rui-Yao Zhang, Mitchell A Pallett, Jamie French, Hongwei Ren, and Geoffrey L Smith. Vaccinia virus btb-kelch proteins c2 and f3 inhibit nf-$\kappa$b activation. *Journal of General Virology*, 103(10):001786, 2022.
